# The super-healing MRL strain promotes muscle growth in muscular dystrophy through a regenerative extracellular matrix

**DOI:** 10.1101/2023.06.29.547098

**Authors:** Joseph G. O’Brien, Alexander B. Willis, Ashlee M. Long, Jason Kwon, GaHyun Lee, Frank Li, Patrick G.T. Page, Andy H. Vo, Michele Hadhazy, Rachelle H. Crosbie, Alexis R. Demonbreun, Elizabeth M. McNally

**Affiliations:** Center for Genetic Medicine, Northwestern University Feinberg School of Medicine, Chicago, IL 60611, USA; Department of Integrative Biology and Physiology, UCLA, Los Angeles, CA; Department of Neurology David Geffen School of Medicine, UCLA, Los Angeles, CA; Department of Pharmacology, Northwestern University Feinberg School of Medicine, Chicago, IL 60611, USA

**Keywords:** muscle, Limb girdle muscular dystrophy, LGMD, TGF-β, super-healing, MRL, extracellular matrix

## Abstract

Genetic background shifts the severity of muscular dystrophy. In mice, the DBA/2J strain confers a more severe muscular dystrophy phenotype, whereas the Murphy’s Roth Large (MRL) strain has “super-healing” properties that reduce fibrosis. A comparative analysis of the *Sgcg* null model of Limb Girdle Muscular Dystrophy in the DBA/2J versus MRL strain showed the MRL background was associated with greater myofiber regeneration and reduced structural degradation of muscle. Transcriptomic profiling of dystrophic muscle in the DBA/2J and MRL strains indicated strain-dependent expression of the extracellular matrix (ECM) and TGF-β signaling genes. To investigate the MRL ECM, cellular components were removed from dystrophic muscle sections to generate decellularized “myoscaffolds”. Decellularized myoscaffolds from dystrophic mice in the protective MRL strain had significantly less deposition of collagen and matrix-bound TGF-β1 and TGF-β3 throughout the matrix, and dystrophic myoscaffolds from the MRL background were enriched in myokines. C2C12 myoblasts were seeded onto decellularized matrices from *Sgcg*^−/−^ MRL and *Sgcg*^−/−^ DBA/2J matrices. Acellular myoscaffolds from the dystrophic MRL background induced myoblast differentiation and growth compared to dystrophic myoscaffolds from the DBA/2J matrices. These studies establish that the MRL background also generates its effect through a highly regenerative ECM, which is active even in muscular dystrophy.

**Brief Summary:** The extracellular matrix of the super-healing MRL mouse strain harbors regenerative myokines that improve skeletal muscle growth and function in muscular dystrophy.

**Graphical Abstract:** 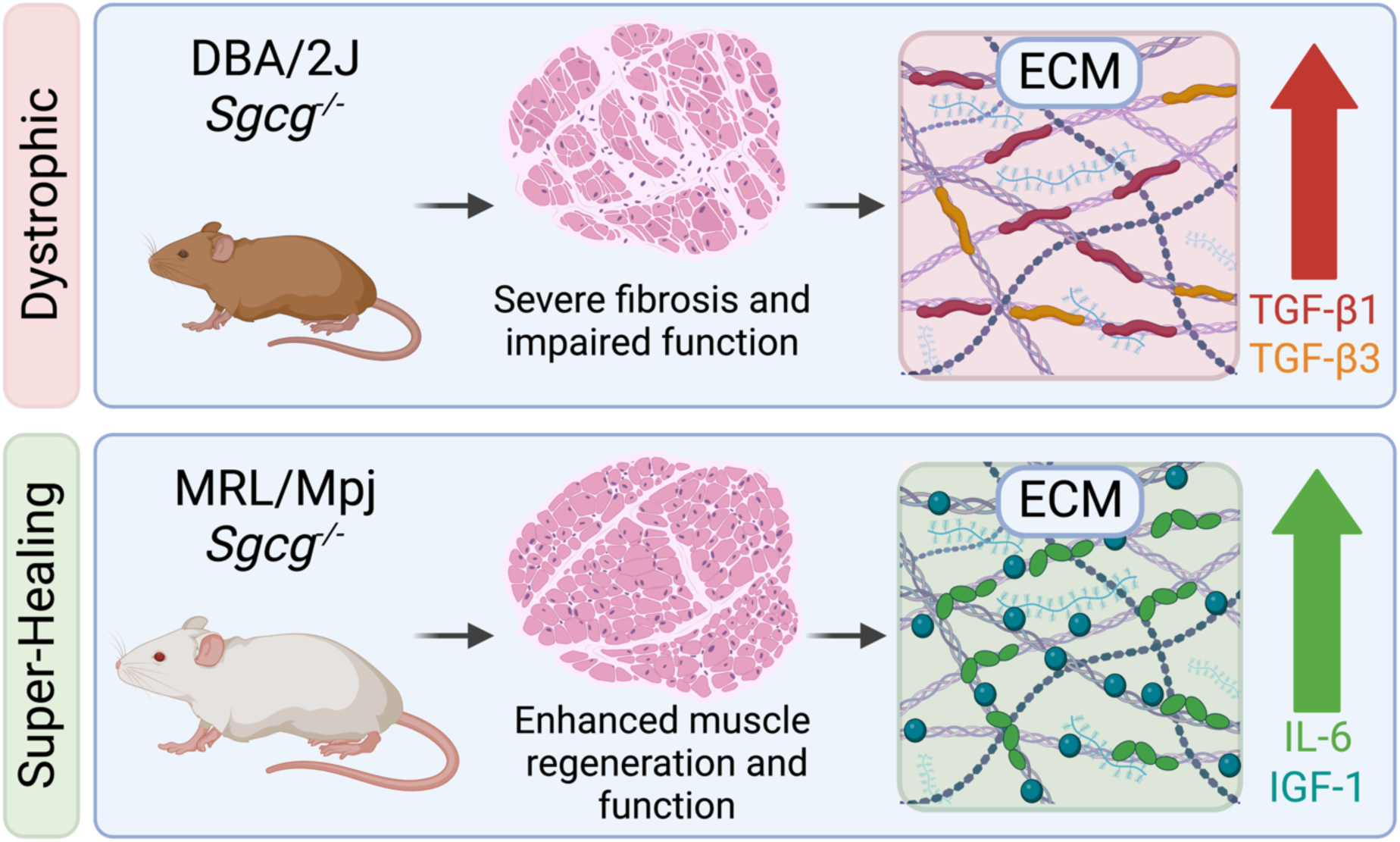

## INTRODUCTION

Genetic disruption of the Dystrophin Glycoprotein Complex (DGC) contributes significantly to muscular dystrophy in humans, and multiple mouse models bear mutations in DGC-encoding genes (1, 2). A shared molecular pathology in these disorders is the interrupted connection of the intracellular cytoskeleton from the plasma membrane and the extracellular matrix (ECM) (3–5). Loss-of-function mutations that disrupt the sarcoglycan subunits within the DGC lead to Limb Girdle Muscular Dystrophy (LGMD) (6). The severity of LGMD caused by mutations in *SGCG*, the gene encoding the γ-sarcoglycan subunit, is highly variable (7). This phenotypic variability has been recapitulated with the generation of *Sgcg* null mice (*Sgcg^−/−^*) and the introduction of this allele into multiple different background strains (8, 9). The DBA/2J (D2) strain intensifies many of the features of muscular dystrophy, including muscle plasma membrane instability, fibrosis, and functional decline (9, 10). Conversely, the genetic backgrounds of C57BL/6 and 129S1/SvlmJ substrains are more resistant to sarcolemma degradation and fibrosis providing partial protection (9, 11, 12).

The Murphy Roths Large (MRL) mouse strain has been studied for its ability to suppress fibrosis and accelerate wound healing in many physiological settings (13, 14). First discovered for its ability to form “scarless” wound healing, the MRL strain is thought to impart both enhanced regeneration and repair (13, 15). In muscle injury, a comparative analysis of the MRL and C57BL/6J (B6) strains after cardiotoxin injury showed rapid reduction of wound size and progenitor muscle cell activation in the MRL strain (16). Enhanced recovery after injury was accompanied by increased expression of antioxidants in the MRL background (16). It was also shown that muscle-derived stem cells isolated from the MRL strain have enhanced expression of *Myod1*, *Myog* and *Pax7* transcripts in response to injury, and hypoxia inducible factor-1 alpha (HIF-1α) contributed to the increase in *Pax7* expression (17).

The ECM mediates muscle regeneration and repair through its structural proteins as well as the smaller cytokine proteins embedded within these structural elements (18, 19). Transforming Growth Factor-β (TGF-β) is increased in both acute and chronic muscle injury, which in turn, further upregulates collagen and fibronectin (20–22). Mouse models of muscular dystrophy in the DBA/2J strain have markedly elevated TGF-β proteins, in part mediated by polymorphisms in the latent TGF-β binding protein 4 (*Ltbp4*) gene as well as a feedback loop with osteopontin/Spp1 (23). The linkage between ECM composition and disease progression has prompted therapeutic development targeting the matrix (24). A recently described method permits evaluation of decellularized dystrophic muscle in an “on slide” format (25). The generation of these “myoscaffolds” leaves a biologically intact ECM which retains cross-sectional spatial information of the dystrophic muscle tissue (25). Using this optimized decellularization method, the ECM protein laminin was observed to undergo remodeling by skeletal muscle progenitor cells in less severely scarred areas of dystrophic muscle, highlighting the importance of specific ECM proteins in mediating cellular regeneration (25).

The MRL background was previously observed to partially mitigate the severity of fibrosis which characterizes *Sgcg*^−/−^ mice in the DBA/2J background (*Sgcg*-D2) (26). The prior breeding strategy evaluated the effect of having a 50% contribution of the MRL background and found the hybrid *Sgcg*-MRL/D2 background demonstrated a significant decrease of fibrosis in skeletal and cardiac muscle (26). The *Sgcg*-MRL/D2 hybrid background showed some evidence for enhanced regeneration with increased embryonic myosin heavy chain expression and a higher proportion of centralized nuclei in several muscle groups compared to the *Sgcg*-D2 background. We now extended these findings by generating the *Sgcg* null allele on a full MRL background, referred to as *Sgcg*-MRL. The histological, functional, and molecular signatures of *Sgcg*-MRL mice were significantly enhanced in comparison to *Sgcg*-D2 mice, demonstrating that the MRL background produces sustained effects over the repeated degeneration that characterizes muscular dystrophy. Prompted by a transcriptomic profile of reduced fibrosis, we interrogated decellularized myoscaffolds to investigate how the super-healing MRL background modified the ECM. We found that C2C12 myoblasts seeded onto the MRL ECM adopted mobility and morphology features of enhanced regeneration. These studies identify the super-healing properties of the MRL background derive significantly from having a regenerative ECM.

## Results

### Reduced muscle fibrosis and enhanced respiratory muscle function in the super-healing MRL background strain

*Sgcg*^−/−^ mice lack the dystrophin-associated protein, γ-sarcoglycan, and this model was previously shown to display a more intense muscular dystrophy process when present in the DBA/2J (D2) background strain, which produced greater muscle fibrosis and correspondingly weaker muscles (9). We queried whether the MRL background could suppress this more severe muscular dystrophy phenotype by backcrossing the *Sgcg*^−/−^ allele into the MRL strain over 10 generations. Backcrosses used both male and female *Sgcg*^+/−^ heterozygous animals. After 10 generations, heterozygous *Sgcg*^+/−^ animals were intercrossed to generate homozygous *Sgcg*^−/−^ animals. The nature of this breeding generates the *Sgcg* exon 2-deleted null allele in the context of the MRL genome.

Terminal evaluation of *Sgcg^−/−^* muscles at 20-weeks of age was used to assess outcome. We compared the respective wildtype (WT) from the MRL and DBA/2J backgrounds (referred to at WT-MRL and WT-D2, respectively) alongside *Sgcg^−/−^* mice from the MRL and DBA/2J backgrounds (referred to as *Sgcg*-MRL and *Sgcg*-D2, respectively). The cohorts were age and sex-matched, with of 5 male and 5 females. Masson Trichrome staining of the diaphragm muscle from 20-week male mice demonstrated markedly reduced fibrosis in *Sgcg*-MRL muscle compared to *Sgcg*-D2 (**Figure 1A**). The *Sgcg*-D2 model had increased accumulation of ECM fibrosis in the diaphragm muscle (**Figure 1B**), with a corresponding decrease in function. S*gcg*-D2 mice demonstrated impaired respiratory function at ten weeks of age as evidenced by elevated Penh, a composite plethysmography measure (27). *Sgcg*-MRL mice had Penh values indistinguishable from WT-MRL mice and WT-D2 mice, consistent with the reduction in fibrosis imparted by the MRL strain improving muscle function (**Figure 1C**). We also evaluated quadriceps muscles and identified significantly reduced pathological features in *Sgcg-*MRL compared to *Sgcg*-D2 (**Figure 1D)**. Fibrosis was reduced in the *Sgcg*-MRL quadriceps muscles compared to *Sgcg*-D2 mice (**Figure 1E**). There were significantly more internal myonuclei in the muscles of the *Sgcg*-MRL (**Figure 1F**), consistent with an enhanced regenerative response in MRL skeletal muscle.

**Figure 1.**
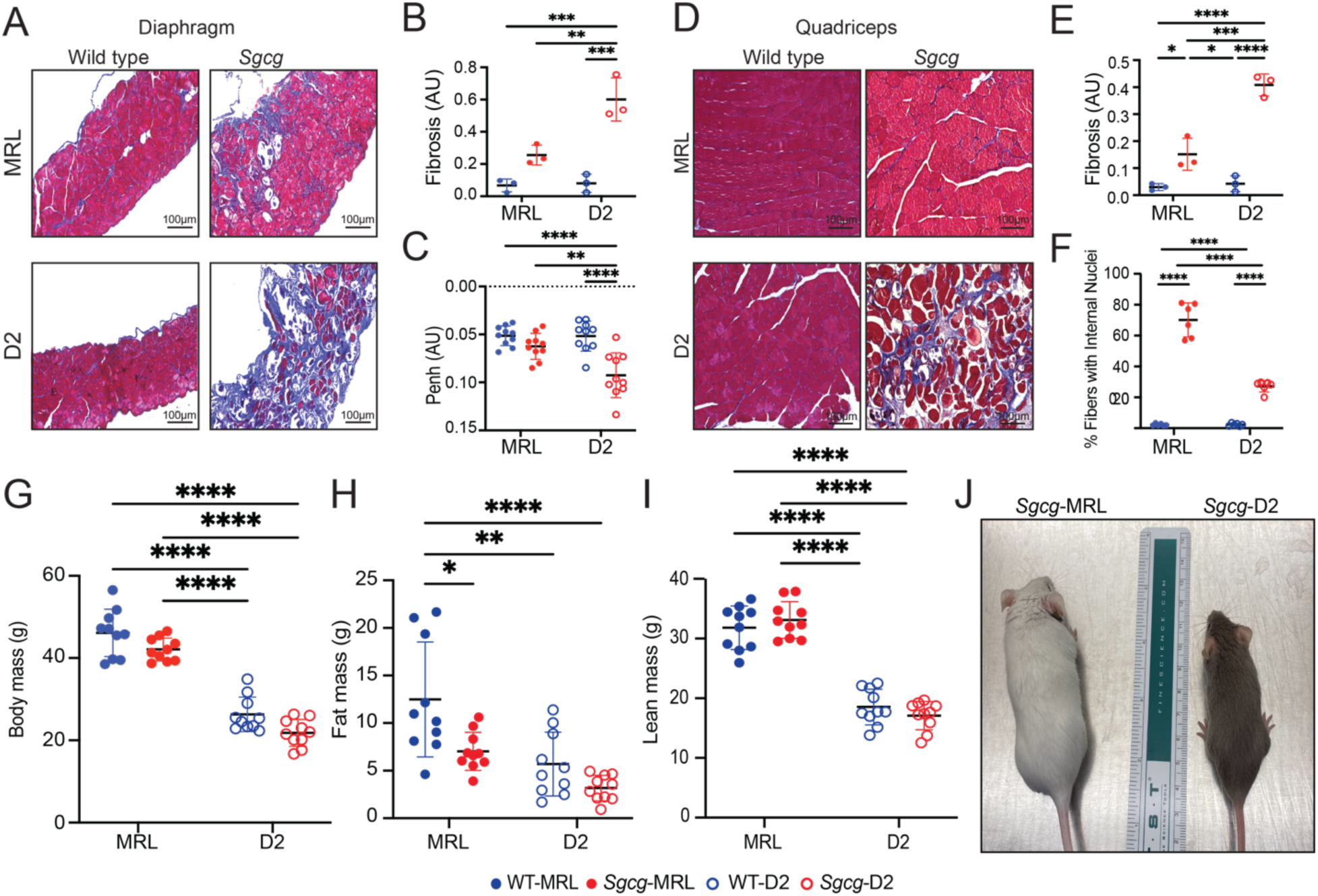
The MRL background reduced fibrosis in the muscles of *Sgcg^−/−^* mice. Comparative phenotypic assessment of skeletal muscle in the *Sgcg*-MRL, *Sgcg*-D2, WT-MRL, and WT-D2 strains was conducted over 20-weeks of cohort including 5 males and 5 females. (**A**) Representative Masson Trichrome staining of the diaphragm muscles (20 weeks) showed the MRL background significantly improved fibrosis in *Sgcg^−/−^* mice. (**B**) Reduction of mean fibrosis (blue staining) relative to total muscle area in the MRL background (WT-MRL 0.03, *Sgcg*-MRL 0.15, WT-D2 0.04, *Sgcg*-D2 0.41 AU). (**C**) Average Penh, a measure of impaired respiratory function, was corrected by the MRL background (WT-MRL 0.05, *Sgcg*-MRL 0.06, WT-D2 0.05, *Sgcg*-D2 0.09 AU). (**D**) Representative images of the quadriceps muscles with reduced fibrosis in *Sgcg*-MRL mice. (**E**) Reduction of mean fibrosis in the quadriceps muscles relative to total muscle area in the MRL strain (WT-MRL 0.07, *Sgcg*-MRL 0.26, WT-D2 0.08, *Sgcg*-D2 0.62 AU). (**F**) Mean percentage of fibers containing centralized nuclei was increased in the *Sgcg*-MRL subjects (WT-MRL 2%, *Sgcg*-MRL 70%, WT-D2 2%, *Sgcg*-D2 27%). (**G**) Body mass was increased in the MRL background. (**H**) Mean fat mass was highest in the WT-MRL cohort (WT-MRL 12.5, *Sgcg*-MRL 7, WT-D2 5.7, *Sgcg*-D2 3.2g). (**I**) Mean lean mass was significantly higher in both MRL models compared to the D2 models (WT-MRL 31.8, *Sgcg*-MRL 18.5, WT-D2 33.1, *Sgcg*-D2 17.1g). (**J**) Representative image depicting the size variation between the *Sgcg*-MRL and *Sgcg*-D2 mice. Graphical quantification of mean ± SD. Two-way ANOVA test with Tukey’s multiple comparison was used to determine statistical significance. * p<0.05, ** p<0.01, *** p<0.001, **** p<0.0001.

Body mass was recorded at 10, 15 and 20 weeks of age and showed a strong strain dependency, as WT-MRL mice were much larger than WT-D2 mice (**Figure 1G**). Body composition analysis by NMR assessed fat and lean contributions of body mass (**Figure 1H and I**, respectively). In the MRL strain, there were sex-specific differences in body mass composition. Although total body mass was similar between males and females WT-MRL mice (**Supplemental Figure 1A**), female WT-MRL mice had significantly increased fat mass and correspondingly reduced lean mass compared to male WT-MRL mice (**Supplemental Figure 1B and C**). This sex-specific body mass composition was absent in *Sgcg*-MRL mice, which exhibited similar fat to lean mass ratios between male and female mice with muscular dystrophy. In the D2 cohorts, both WT-D2 and *Sgcg*-D2 lacked any detectable differences in total body mass and body mass composition (**Supplemental Figure 1D**). By 20 weeks of age, a stage when *Sgcg*-D2 mice display kyphoscoliosis and wasting in both male and female mice, *Sgcg-*MRL mice appeared outwardly indistinguishable from their WT counterparts **(Figure 1J).**

### Larger muscles in MRL mice compared to D2 mice

The increase in centrally-nucleated myofibers in *Sgcg*-MRL muscle compared to *Sgcg*-D2 muscle suggests the MRL background promotes growth and regeneration. To further investigate muscle morphometric features, we evaluated the *Tibialis anterior* (TA) muscles as these muscles are also suitable for physiological assessment. Laminin-α2 (LAMA2) staining was used to outline myofibers and assess fiber size and myofiber cross sectional area (**Figure 2AB**). A higher proportion of smaller myofibers was present in the TA muscles from *Sgcg*-D2 mice, and these small myofibers were not apparent in TA muscles from *Sgcg*-MRL mice. Further analyses of the cross-sectional area (CSA) and minimum Feret diameter demonstrated a decrease in muscle fiber size in the *Sgcg*-D2 mice compared to fibers from WT-D2 and *Sgcg*-MRL mice (**Figure 2C and D**), and these findings are consistent with the impaired growth of *Sgcg*-D2 muscle compared to the pro-regenerative environment of the MRL background strain. Fiber type composition in the TA muscles from the MRL mouse strain contained a higher proportion of fast twitch (type IIB) muscle fibers compared to D2 muscles, and this was apparent in both the WT-MRL and *Sgc*g-MRL mice (**Supplemental Figure 2A**). These results were further validated by fiber counting cryosections, which confirmed the higher proportion of type IIB fibers in the MRL background (**Supplemental Figure 2B**).

**Figure 2.**
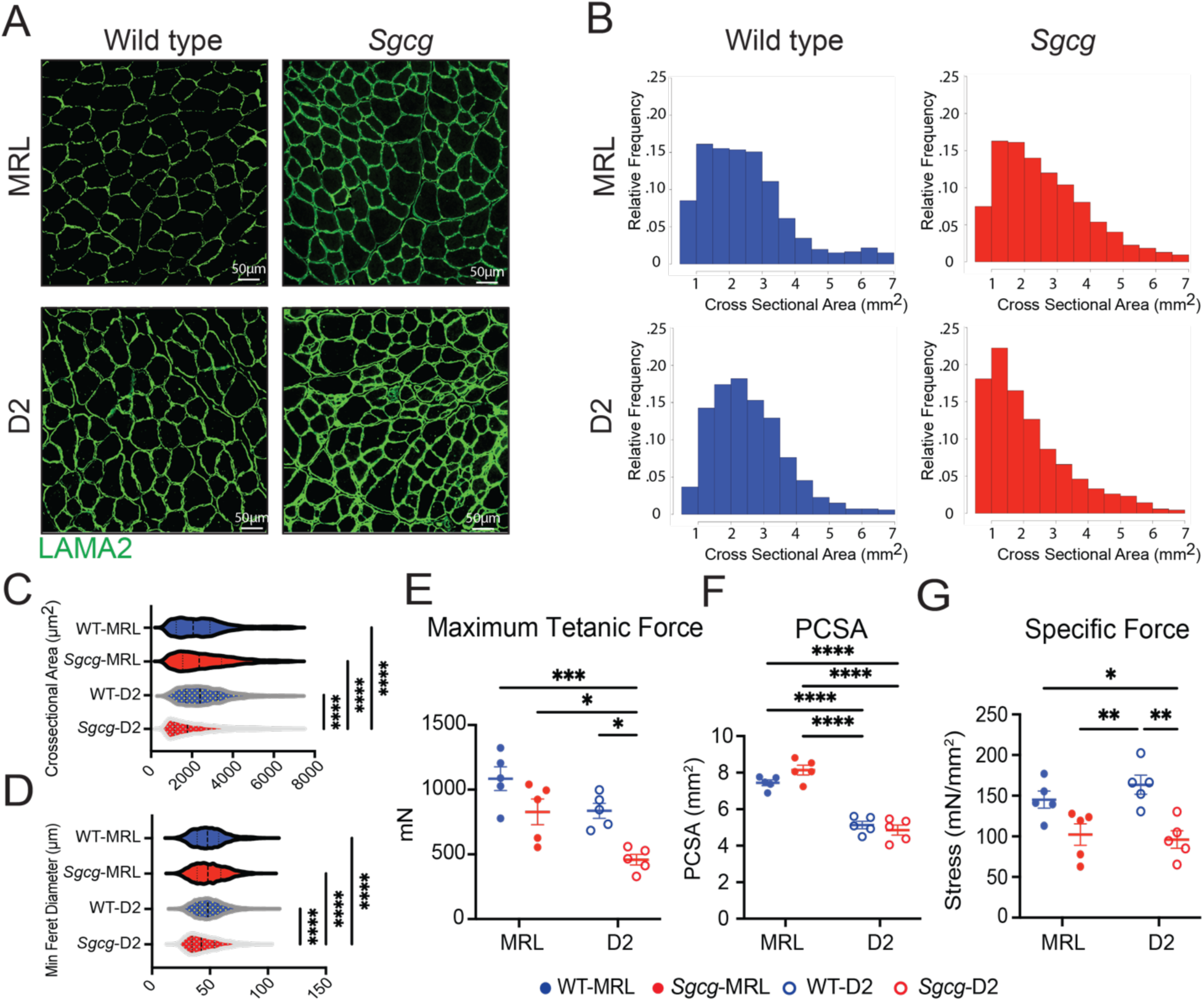
The MRL strain increased myofiber size and maximum tetanic force in *Sgcg*^−/−^ mice. TA muscle was analyzed at 20 weeks. (**A**) Representative immunofluorescence microscopy (IFM) images showed larger myofibers in the TA of the *Sgcg*-MRL stained with laminin-α2 (LAMA2). (**B**) Cross sectional area (CSA) of myofibers was shifted rightward in *Sgcg-*MRL muscle. (**C**) CSA was increased in the MRL background (WT-MRL 2544, *Sgcg*-MRL 2619, WT-D2 2560, *Sgcg*-D2 2154 μm^2^). (**D**) MRL background increased the Minimum Feret Diameter in the *Sgcg^−/−^* muscle (WT-MRL 49.1, *Sgcg*-MRL 49.6, WT-D2 49.6, *Sgcg*-D2 45.1μm). (**E**) Force measurements from male TA muscles showing maximum tetanic force was significantly reduced in the *Sgcg*-D2 strains compared to all other strains (WT-MRL 1085, *Sgcg*-MRL 828.5, WT-D2 836.3, *Sgcg*-D2 458.7mN). (**F**) Physiological Cross-Sectional Area (PCSA) was smaller in D2 compared to MRL muscle for both *Sgcg^−/−^* and WT (WT-MRL 7.44, *Sgcg*-MRL 8.14, WT-D2 5.13, *Sgcg*-D2 4.85mm^2^). (**G**) Specific force was reduced for *Sgcg^−/−^* compared to WT in the D2 strain but not the MRL strain (WT-MRL 145, *Sgcg*-MRL 102.3, WT-D2 163.5, *Sgcg*-D2 96.0mN/mm^2^). Graphical quantification of mean ± SD. Two-way ANOVA test with Tukey’s multiple comparison was used to determine statistical significance. * p<0.05, ** p<0.01, *** p<0. 001, **** p< 0.0001.

To evaluate muscle force production, *in-situ* force analysis was performed on the TA muscle. Male *Sgcg-*MRL TA muscle produced twice the maximum tetanic force compared to male *Sgcg*-D2 TA muscle (**Figure 2E**). This difference was not evident in female mice due high variability in muscle force measurements **(Supplemental Figure 3A**). The physiological cross-sectional area (PCSA) of TA muscles was larger for MRL mice compared to D2 mice for both males and females, consistent with the larger size of MRL compared to D2 mice (**Figure 2F and Supplemental Figure 3B**). In both male and female mice, specific force was similar between the *Sgcg*-MRL and *Sgcg*-D2 muscles, consistent with MRL muscles generating strength proportional to their larger size.

### Upregulation of regenerative pathways in Sgcg-MRL muscles and increased inflammatory gene expression in Sgcg-D2 muscles

Bulk RNA sequencing was performed on quadriceps muscles harvested from 16-week-old WT-MRL, *Sgcg*-MRL, WT-D2 and *Sgcg*-D2 mice. Principal Component Analysis of the RNAseq dataset revealed divergent clustering with effects of both genetic strain and the *Sgcg*^−/−^ disease mutation (**Supplemental Figure 4**). Direct comparison of the *Sgcg*-MRL and *Sgcg*-D2 muscles revealed the MRL background resulted in a greater expression of genes associated with muscle development and differentiation, while *Sgcg-*D2 tissue expressed genes linked to inflammatory responses, extracellular matrix development and TGF-β signaling (**Figure 3A**). Representitive heatmaps demonstrate an upregulation of gene expression associated with ECM and the TGF-β signaling pathway were upregulated in the D2 model (**Figures 3B and 3C**). The full, clustered list of these genes are presented in (**Supplemental Figure 5 and 6**). *TGFB1* was upregulated in *Sgcg*-D2 and downregulated in *Sgcg*-MRL (**Figure 3D**). TGF-β signaling was evaluated in quadriceps muscle by immunoblot analysis and showed excess phosphorylated SMAD3 (pSMAD3) in *Sgcg*-D2 muscle compared to *Sgcg*-MRL (**Figure 3E and F**).

**Figure 3.**
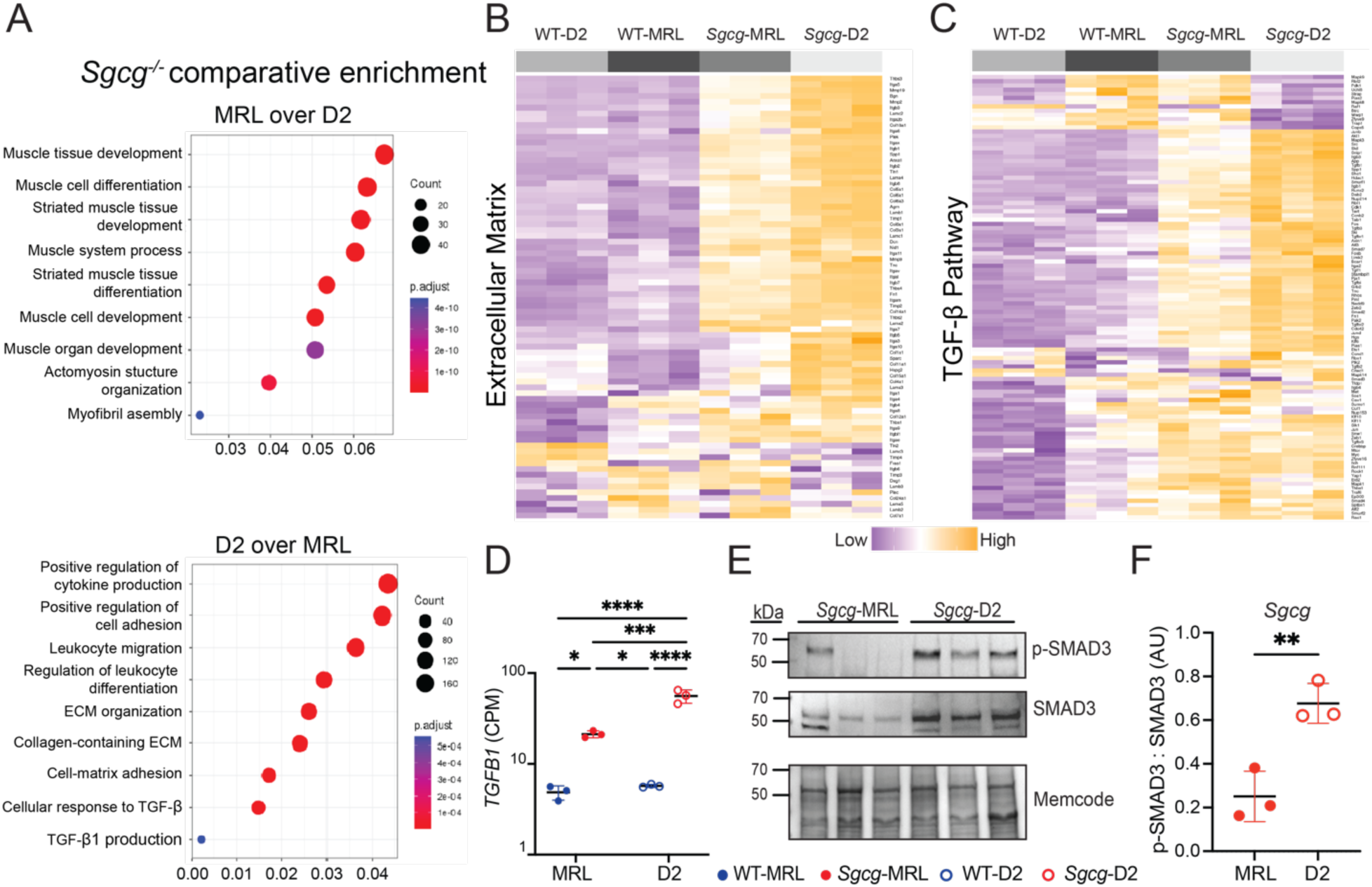
TGF-β gene expression and signaling were downregulated in *Sgcg*-MRL skeletal muscle compared to *Sgcg*-D2 muscle. RNA sequencing was performed on quadriceps muscles from 16-week-old female mice in biological triplicate. Gene expression was normalized to Counts Per Million (CPM). (**A**) A comparison of *Sgcg*-MRL versus *Sgcg*-D2 transcriptomic profiles showed muscle development and differentiation genes to be highly enriched the *Sgcg*-MRL background (34). In contrast, immune response, extracellular matrix, and TGF-β pathways were highly enriched in the *Sgcg*-D2 cohort. (**B**) Clustered heatmap shows reduction of extracellular matrix genes in *Sgcg*-MRL muscle. (**C**) Clustered heatmap shows reduction in TGF-β pathway genes in *Sgcg*-MRL muscle. (**D**) Comparative log scale analysis showed substantially higher average expression of *TGFB1* in the *Sgcg*-D2 cohort (WT-MRL 4.86, *Sgcg*-MRL 21.3, WT-D2 5.73, *Sgcg*-D2 55.9 CPM). (**E**) Immunoblotting of quadriceps muscles from *Sgcg*-D2 and *Sgcg*-MRL mice showed reduced phosphorylated SMAD3 (p-SMAD3) and total SMAD3 in *Sgcg*-MRL. (**F**) The ratio of p-SMAD3 to total SMAD3 was quantified and indicated TGF-β signaling was decreased in the MRL background. Graphical quantification of mean ± SD. Two-way ANOVA was used to determine statistical significance in 4B, and Student’s t-test was used in 4D. * p<0.05, ** p<0.01, *** p<0.001, **** p<0.0001.

### The MRL strain has a regenerative extracellular matrix

The molecular composition of the extracellular matrix was investigated in decellularized extracellular matrices (dECM) prepared from *Sgcg*-MRL and *Sgcg*-D2 quadriceps muscles. This “on slide” decellularization protocol was optimized to generate myoscaffolds following the methods in (28, 29). *Sgcg*-MRL and *Sgcg*-D2 quadriceps sections were incubated in 1% SDS and subsequently stained with Hematoxylin & Eosin (H&E) and Sirius Red to confirm removal of cellular material and retention of an intact ECM (**Figure 4A and B**, respectively). Representative images of dECMs confirm removal of intracellular components compared to control sections treated with 0% SDS which retained the cellular components and validated the effectiveness of this decellularization method. *Sgcg*-MRL myoscaffolds displayed reduced collagen, evidenced through Sirius Red, compared to *Sgcg*-D2 myoscaffolds, consistent with the antifibrotic role of the MRL background in *Sgcg*-MRL muscle.

**Figure 4.**
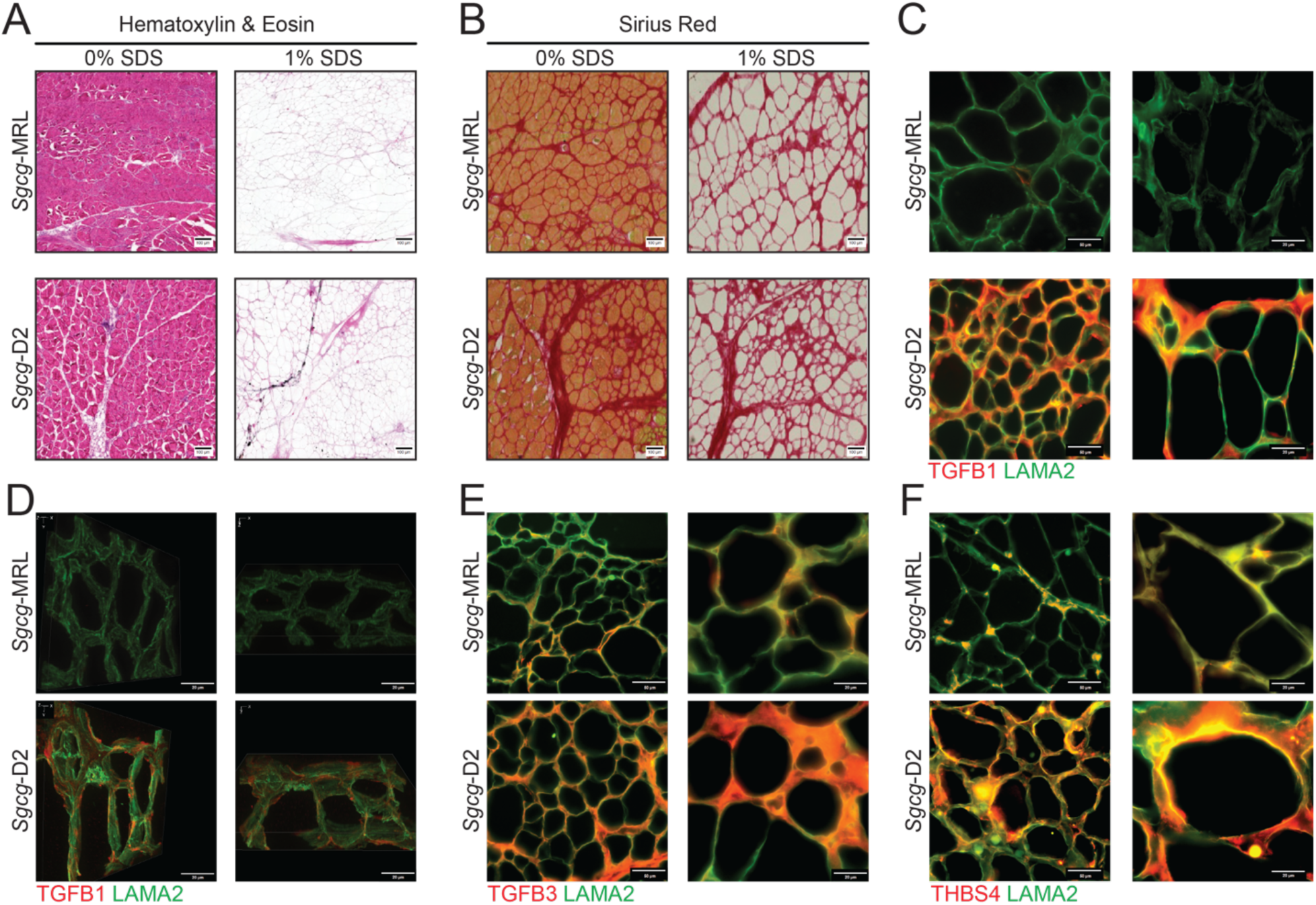
Reduction of TGF-β proteins in decellularized matrices from *Sgcg*-MRL muscle compared to *Sgcg*-D2 muscles. Decellularized extracellular (dECM) myoscaffolds were generated from *Sgcg*-MRL and *Sgcg*-D2 quadriceps muscles. Myoscaffold sections were stained with (**A**) Hematoxylin & Eosin and (**B**) Sirius Red demonstrating removal of cellular content following detergent treatment (1.0% SDS) with maintenance of matrix components. Control samples taken from adjacent sections were incubated in PBS only (0%SDS). (**C**) Representative IFM images of dECM myoscaffolds co-stained with antibodies to detect TGF-β1 (red) and LAMA2 (green). Extracellular TGF-β1 protein expression was dramatically reduced in the MRL background. (**D**) 100x z-stack images of TGF-β1 (red) and LAMA2 (green) demonstrate excess TGF-β1 distributed throughout the matrix in the dystrophic D2 model. (**E)** Representative IFM images of dECM sections, co-stained with antibodies to detect TGF-β3 (red) and LAMA2 (green) showing reduction of TGF-β3 in *Sgcg*-MRL myoscaffolds. (**F**) dECM myoscaffolds from *Sgcg*-MRL (34) and *Sgcg*-D2 (bottom) showed reduced thrombospondin-4 (THBS4, red) and LAMA2 (green) in the *Sgcg*-MRL muscle.

Representative images of dECM myoscaffolds co-stained with anti-TGF-β1 and anti-LAMA2 exhibited reduced TGF-β1 signal in the dECM of the *Sgcg*-MRL model (**Figure 4C**). Further visualization through 3D z-stack imaging demonstrated minimal TGF-β1 in *Sgcg*-MRL scaffolds, while TGF-β1 signal was abundant throughout the matrix of *Sgcg*-D2 mice (**Figure 4D**). Similarly, anti-TGF-β3 and anti-LAMA2 co-staining of the myoscaffolds revealed that TGF-β3 was also reduced by the MRL background (**Figure 4E**). Thrombospondin-4 (THSB4), a multifunctional glycoprotein that modifies muscular dystrophy outcome, was also markedly downregulated in the *Sgcg*-MRL myoscaffolds but was abundant in the *Sgcg*-D2 dECM (**Figure 4F**). Together, these results indicate a dramatic decrease in TGF-β signaling induced by the MRL background, which in turn, likely promotes the pro-regenerative nature of the MRL background.

### Sgcg-MRL dECM myoscaffold enhanced differentiation in seeded C2C12 myoblasts

The extracellular environment of skeletal muscle provides an essential substrate to support regeneration and formation of myotubes after injury (30, 31). Given the differential protein deposition of the dECMs of the *Sgcg*-MRL and *Sgcg*-D2 muscles, we investigated how these matrices altered motility and differentiation of myoblasts. C2C12 myoblasts were seeded on dECM myoscaffolds. Following a five-day differentiation protocol, the slides were fixed and stained with antibodies to desmin (DES) and LAMA2 (**Figure 5A and B**). Desmin-positive cells on the *Sgcg*-MRL myoscaffolds were larger and covered more area compared to myoblasts similarly seeded on *Sgcg*-D2 myoscaffolds. On the *Sgcg*-D2 scaffolds, desmin-positive cells were small with little cellular area, consistent with poor differentiation (**Figure 5B**). Desmin expression and cell morphology were quantified using Image J software from six representative images captured from three animals of each genotype. When standardized to the number of nuclei, the percent area of desmin fluorescence was significantly increased in myoblasts seeded on the *Sgcg*-MRL myoscaffolds compared to those seeded on the *Sgcg*-D2 myoscaffolds (**Figure 5C**). Myoscaffold strain background had a significant influence on the average cell size, circularity, and perimeter (**Figure 5D, E and F, respectively**), where MRL-seeded myoblasts exhibited an increase in cell size and had a less circular morphology. Similarly, myoblasts seeded on MRL scaffolds exhibited a significant increase in maximum and minimum Feret diameter (**Figure 5G and H**). These measures indicate that the dECM myoscaffolds of the MRL strain promoted greater growth and development of myoblasts.

**Figure 5.**
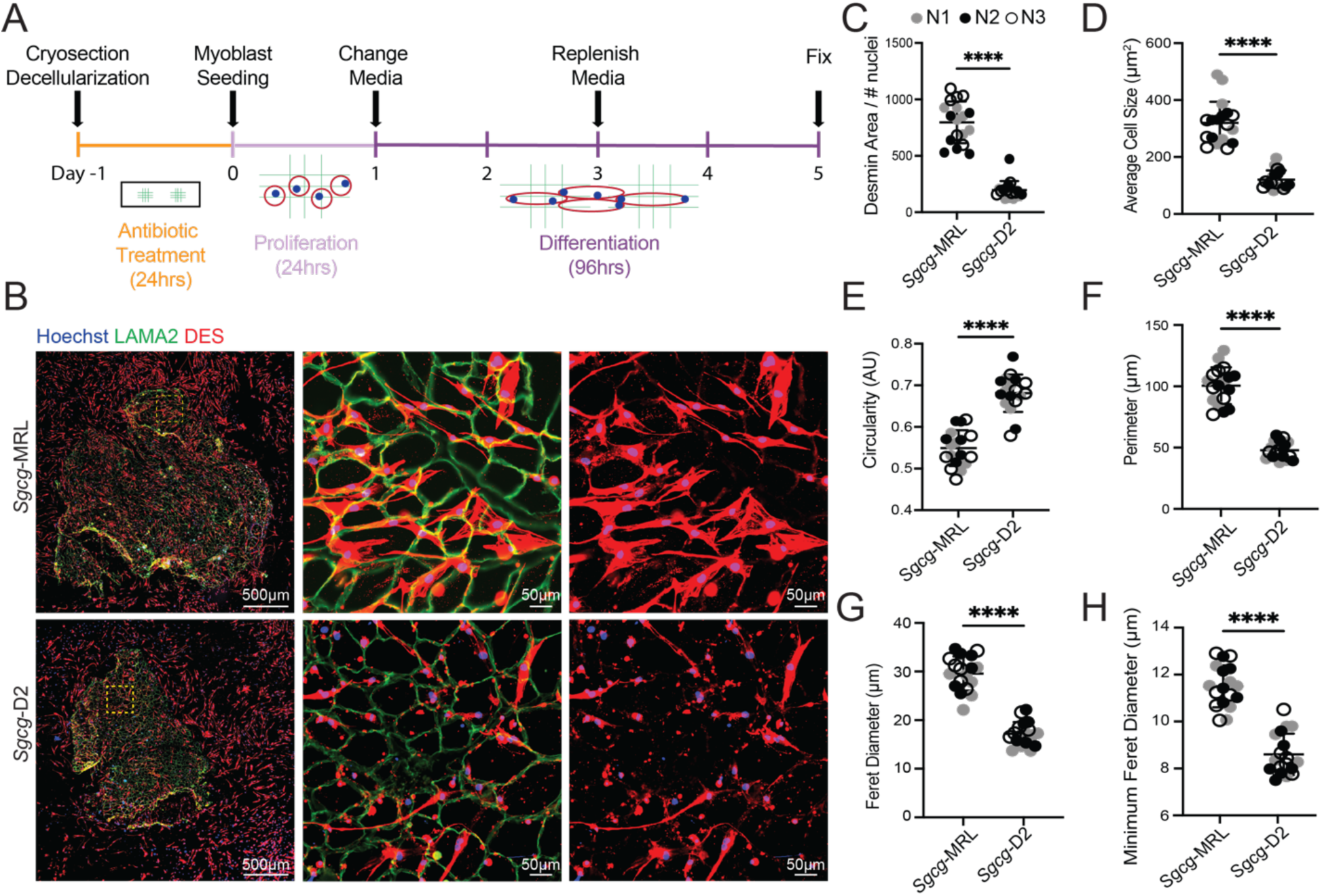
Decellularized myoscaffolds from *Sgcg*-MRL muscle promote growth and differentiation of C2C12 myoblasts. (**A**) C2C12 myoblasts were seeded and proliferated on dECM myoscaffolds from *Sgcg*-MRL (34) and *Sgcg*-D2 (bottom) for 24 hours. Then growth media was replaced with low-serum differentiation media for 96 hours. Slides were fixed with PFA for IFM. (**B**) Imaging of dECM myoscaffolds seeded with C2C12 myoblasts. The left column represents entire myoscaffolds with seeded myoblasts co-stained with LAMA2 (green), desmin (DES, red), and Hoechst (blue) to stain nuclei. The yellow region of interest is magnified in the middle and right columns. DES (red) staining in the right column shows myoblast morphology. (**C-H**) Three independent biological replicates of the *Sgcg*-MRL and *Sgcg*-D2 dECMs were analyzed (labelled as N1 = gray, N2 = black, N3 = white). For each sample, six images of myoblasts were captured at 20x magnification. **C** shows the percentage area of desmin positivity relative to the number of nuclei in the field (*Sgcg*-MRL 797.0, *Sgcg*-D2 197.4). **D** shows the average cell size (*Sgcg*-MRL 319.7, *Sgcg*-D2 120.8 mm^2^). **E** shows circularity (*Sgcg*-MRL .549, *Sgcg*-D2 .681 AU). **F** shows myoblast perimeter (*Sgcg*-MRL 100.5, *Sgcg*-D2 47.8mm). **G** shows Feret diameter (*Sgcg*-MRL 29.6, *Sgcg*-D2 17.2mm). **H** shows minimum Feret diameter (*Sgcg*-MRL 11.5, *Sgcg*-D2 8.6mm). Graphical quantification of mean ± SD. A Student’s t-test was used to determine significance. **** p<0.0001

### MRL background has increased circulating growth factors

The capacity of skeletal muscle to repair and regenerate after injury is directly influenced by circulating cytokines and growth factors (32, 33). We used the SomaScan aptamer assay to assess the proteomic profile of serum collected from the *Sgcg*-MRL and *Sgcg*-D2 cohorts at 20 weeks of age. Comparative enrichment analysis between the models found the MRL serum to harbor increased expression of proteins associated with growth factor activity and wound healing (**Figure 6A**). Heat maps depicting differentially expressed proteins (FDR p-value <0.1) associated with growth factor activity (34) and wound healing (bottom) indicate upregulation in the MRL strain, where the samples are ordered by sex and significance. IGF-1 and IL-6 were among the most differentially expressed proteins in both pathways (**Figure 6B**). Enzyme-linked immunosorbent assays (ELISAs) were conducted in the same serum samples for IL-6 (**Figure 6C**) and IGF-1 (**Figure 6D**). Both proteins were significantly upregulated in the MRL background, consistent with the SomaScan results. We further profiled the expression of these two proteins within decellularized myoscaffolds from dystrophic muscle in the MRL and D2 backgrounds and found the *Sgcg*-MRL myoscaffolds had visible deposition of both IL-6 and IGF-1 in their matrices, unlike *Sgcg*-D2 myoscaffolds (**Figure 6E** and **F**). These data highlight that the pro-regenerative milieu of the MRL muscle is also reflected in the serum protein profiles.

**Figure 6.**
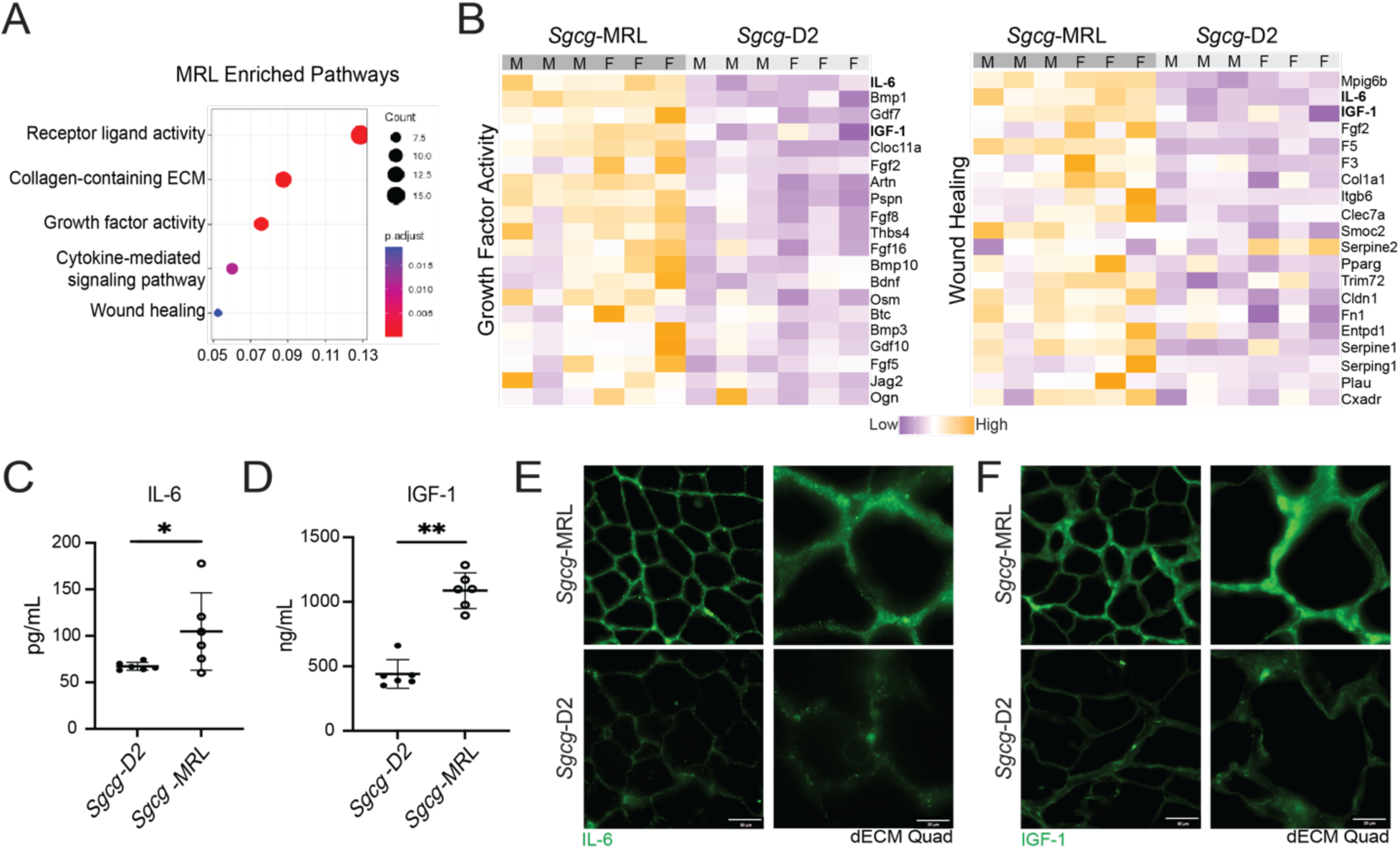
IGF-1 and IL-6 were upregulated in *Sgcg*-MRL serum. Serum was collected from *Sgcg*-MRL and *Sgcg*-D2 cohorts at 20 weeks of age and evaluate using the SOMAscan aptamer assay. (**A**) Pathway enrichment analysis showed receptor ligand, collagen containing ECM and growth factor activity highly enriched in *Sgcg*-MRL serum. (**B**) Heatmaps of proteins in the growth factor activity (left) and wound healing (right) pathways upregulated in *Sgcg*-MRL mice. IL-6 and IGF-1 (bold) were among the most differentially expressed proteins in both pathways. IL-6 (**C**) and IGF-1 (**D**) upregulation was verified using ELISA analysis. (**E**) and (**F**) show matrix deposition of IL-6 and IGF-1 increased in decellularized myoscaffolds from *Sgcg*-MRL but not *Sgcg*-D2 muscles. Graphical quantification of mean ± SD. Statistical significance was determined using the Kolmogorov-Smirnov test. * p<0.05, ** p<0.01

## DISCUSSION

### The MRL background promotes sustained muscle regeneration in muscular dystrophy

The enhanced wound healing properties of the MRL strain have been described after many different forms of acute injury like earhole punching or acute injury of cartilage or skin (13, 15). Many these studies evaluating the wound healing properties of the MRL background are conducted on younger animals, since with age MRL model is susceptible to autoimmune disorders (35, 36). We focused our study on the first 20 weeks of age, finding evidence for a sustained contribution of the MRL background to improve the severe fibrosis characteristic of the dystrophic *Sgcg* null allele. In mice and humans with muscular dystrophy, the diaphragm and respiratory muscles are highly impaired (37, 38), and so it is notable that the MRL background was able to correct functional deficits in the dystrophic mouse model. The protective qualities of the MRL background were not restricted to just the diaphragm muscle since the other studied muscles also showed functional and histological improvement, including marked reduction of fibrosis and enhanced features of muscle regeneration. Muscular dystrophy preferentially degrades fast twitch myofibers (Type IIB), causing a fiber type shift from fast to slow twitch (Type I) (39, 40). The MRL background protected against this fiber type shift, where a higher proportion of MYH4 positive fibers (Type IIB) were present, and MYH7 positive fibers (Type I) were almost completely absent. It is possible over even longer periods of time that the MRL background may begin to uncover some of the autoimmune features in this background.

### Sex influences body mass composition in the MRL strain

MRL mice are larger than many other mouse strains, nearly twice the size of other mice, and along with this, the MRL strain carries a distinct metabolic profile (16, 41, 42). Embryonic features of the metabolism are retained longer in MRL mice, and these features have also been associated with decreased reactive oxygen species (ROS), and consequent stem cell activation and tissue regeneration (41). MRL mitochondria contain two naturally-encoded heteroplasmies, which have been found to decrease severity of muscle disease (43). It has been shown that MRL mice adapted to a high fat diet by metabolically shifting from carbohydrate metabolism toward β-oxidation. In this setting of high fat diet, MRL still experienced significant weight gain, but this metabolic shift protected against cardiac hypertrophy, hyperglycemia, and insulin resistance (42). Although we observed no difference in overall body mass between males and females WT-MRL mice, the body mass composition varied considerably with females gaining significantly more fat mass than their male counterparts. This sexually dimorphic pattern of weight gain was not apparent in the *Sgcg*-MRL or D2 cohorts, where body composition was similar between males and females, likely because the dystrophic phenotype was dominating body composition.

### Suppression of TGF-β signaling in the MRL background

The TGF-β proteins (β1, β2, β3) promote fibrosis in many settings, including muscular dystrophy (21, 44, 45). In part, TGF-β induced fibrosis is dictated by genetic modifiers like *LTBP4* and *SPP1* (23, 46)*. LTBP4* suppresses active TGF-β in dystrophic muscle. Polymorphisms of *LTBP4* in the D2 background are associated with increased TGF-β signaling through SMAD2/3 phosphorylation (46, 47). The activation of SMAD2/3 upregulates gene expression ECM associated genes (48), and the DBA/2J strain carries an allele of *Ltbp4* that enhances active TGF-β, contributing significantly to the exacerbation of muscular dystrophy by the DBA/2J background (12). We demonstrate that TGF-β signaling is suppressed, with SMAD2/3 phosphorylation in the MRL background. This reduction of TGF-β signaling corresponded with enhanced cell growth features in myoblasts seeded onto the MRL-dECM. These findings are consistent with other injury settings, including cartilage (49), where downregulation of TGF-β signaling strongly correlated with improved cartilage repair.

Decellularized ECM scaffolds are useful platforms to study matrisomal protein composition and tissue regeneration (50–52). We employed the decellularization methods developed by (25) and found the highly fibrotic *Sgcg*-D2 muscles were characterized with excess TGF-β1 deposited throughout the matrix. In contrast, the MRL background had little to no TGF-β1 in its matrix. Similarly, expression of extracellular matrix proteins TGF-β3 and THBS4 was notably diminished in the MRL background. THBS4 is a key protein in the assembly of the matrix after injury and promotes tendon strength and repair (53). Analysis of human muscle biopsies from human muscular dystrophy patients found enriched THBS4 expression, consistent with dystrophic mouse models (54, 55). The reduction of THBS4 and TGF-β proteins in MRL matrix corresponds to the decreased collagen accumulation and is consistent with markedly delayed progression of the muscular dystrophy process.

### The MRL harbors a regenerative matrix that promotes muscle growth

Compositional components of the matrix can provide instructive cues during muscle regeneration (30, 56). Muscle regeneration involves a multifaceted, coordinated response which can be interrupted by inflammatory factors (19), including those components present in dystrophic muscle (54, 57). We found the dECM derived from the dystrophic MRL background promoted myoblast shape and size, indicating cell-extrinsic healing properties of the MRL background. We found the serum of *Sgcg*-MRL model to be enriched with proteins associated with growth factor activity and wound healing. While excess TGF-β acts as a potent inhibitor of myoblast differentiation, IGFs stimulate proliferation and regeneration (32). The increased circulation of IGF-1 and IL-6 are positioned to mediate the pro-regenerative qualities observed in the MRL matrix (58–60). IGF-1 is a circulating growth factor that promotes anabolic pathways in skeletal muscle and prevents age-related sarcopenia (58). Overexpression of IGF-1 reduces muscle pathology and stabilizes the sarcolemma in the *mdx* model (61). IL-6 is a secreted glycoprotein that also serves as a myokine, where it is necessary for muscle regeneration (60, 62, 63). Upregulation of IL-6 has been observed in MRL serum, where it has been thought to contribute to autoimmune conditions (64). However, in this setting, the increase in IL-6 is also present as matrix deposition of this myokine, where it can directly contribute to muscle regeneration and the protective, pro-regenerative microenvironment observed in the *Sgcg*-MRL model. This study demonstrates the fundamental influence genetic background has on dystrophic progression and highlights the importance of the extracellular microenvironment in muscle regeneration and repair.

## MATERIALS AND METHODS

### Animals

All animals were bred and housed in a pathogen free facility. Wild-type (WT-D2) mice were from the DBA/2J strain (Jackson Laboratories; Strain # 000671) and (WT-MRL) were from the MRL/Mpj strain (Jackson Laboratories; Strain #000486). The sarcoglycan γ-null (*Sgcg*) mice were previously generated by deleting exon 2 and then breeding more than ten generations into the C57BL/6J background (8). Mice were backcrossed over 10 generation to the DBA/2J (*Sgcg-D2*) and MRL (*Sgcg-MRL*) strains. Male and female mice were used for all experiments, unless otherwise noted. Mice were bred and housed in a specific pathogen free facility on a 12-hour light/dark cycle and fed *ad libitum*. Where possible, physiological analyses were performed blinded to genotype and background strain, although the size/appearance differences between MRL and DBA/2J made blinding to this aspect impossible. Cohorts were randomized to ensure even sex distribution across cohorts and approximate age matching.

### Study Approval

All procedures using mice followed *Guide for the Care and Use of Laboratory Animals* and were approved by Northwestern University’s Institutional Animal Care and Use Committee.

### Histology

Quadriceps muscle was harvested and fixed in 10% formalin. Hematoxylin and eosin (cat# 12013B and 1070C; Newcomer Supply), Masson Trichrome (cat# HT15; Newcomer Supply) and Sirius Red (cat# 24901250; Polysciences, Inc) staining was performed per manufacturer’s protocol. For all genotypes, images were acquired on the Keyence BZ-X810 microscope with the same exposure settings. Masson Trichrome image assessment was performed on representative 20x images from 3 male biological replicates using Fiji (NIH), where the fibrotic content was standardized to full muscle area.

### Body weight and mass composition analysis

Total body mass was determined at 10, 15, and 20 weeks. Fat mass, lean mass and hydration ratio were determined at 20 weeks of age, using the non-invasive NMR method provided by EchoMRI^TM^. Each cohort consisted of 10 subjects, 5 male and 5 female.

### Plethysmography

Unanesthetized whole-body plethysmography (WBP) was performed at 10 weeks using a Data Sciences International, Buxco Finepointe 4-site WBP as described in (65, 66). Chambers were calibrated and then individual mice were placed inside the chamber for a 120-minute acclimation period prior to experimental recording. Data was collected for 10 minutes while mice were in a resting phase. Studies were performed at room temperature. Data points were filtered to include breaths with a 0 Rejection Index (Rinx) and within a frequency range of 100-250 breaths per minute. Penh was normalized to body mass (g).

### Serum collection

Serum was acquired and processed as described (67). Briefly, using a heparinized capillary tube (20-362-566; Fisher Scientific, Waltham, MA) blood was collected by means of retro-orbital puncture into a Microtainer™ Gold Top Serum Separator tube (365967 Becton Dickinson, Franklin Lakes, NJ) and centrifuged at 8,000 x g for 10 min. Serum fractions were collected at 20 weeks of age and stored at −80°C.

### Serum based ELISA analyses for Circulating proteins

Serum fractions from the *Sgcg*-MRL and *Sgcg*-D2 cohorts were collected in 3 male and 3 female subjects at 20 weeks of age. IL-6 and IGF-1 serum levels were assessed using the Mouse IL-6 Quantikine ELISA Kit (R&D Systems, M6000B) and Mouse IGF-1 Quantikine ELISA kit (R&D Systems, MG100), respectively, according to the manufacturers’ instructions.

### Serum aptamer profiling assay and analysis

The SOMAscan^TM^ assay reports 7322 aptamer-based proteomics results per sample in units of Relative Fluorescent Units, which were read into R studio using the SomalogicsIO R package. Linear mixed modeling was performed using the dream package in R. Data were imported and pre-processed, and a linear mixed model was fit to the data using the lme function. Model fit and estimated parameters were obtained using the summary function, and fixed and random effect estimates were extracted using the fixef and ranef functions, respectively. Model diagnostics, including checking the normality of the residuals, visualizing the residuals against the predicted values, and checking for heteroscedasticity and outliers, were performed using the qqmath, plot, and plot functions. Hypothesis tests were conducted using the anova function, and confidence intervals for the estimated parameters were computed using the confint function.

### In-situ force and fatigue

The tibialis anterior (TA) was assayed using an Aurora Scientific 1300A Whole Animal System (Cat #1300A; Aurora Scientific, Aurora, ON, Canada). Mice were anesthetized via isoflurane, and the initial muscle isolation was performed under a higher isoflurane concentration (1L/min of 2.0% isoflurane in 100% O_2_). To prepare the TA for in-situ force measurement, an incision was made in the skin from the anterior side of the left foot up toward the left knee. This allowed for access to both the distal TA tendon and the anterior compartment of the lower hindlimb. Once the fascia was removed, the distal TA tendon was cut, and the TA was isolated via blunt dissection. A braided silk suture (4-0 diameter) was tied around the distal tendon before the mouse was transferred to the experimental setup. The anesthesia was then reduced (1L/min or 1.5% isoflurane in 100% O_2_) to minimize any deleterious effects induced by long term exposure. It was noted during setup that MRL mice required a higher dose of isoflurane (2-2.5% during isolation, 1.8-2% during experimentation). This is likely due to the larger size of the MRL model. The free end of the tendon suture was then tied to the lever arm of a 5N dual-action force transducer (305C-LR; Aurora Scientific, Aurora, ON, Canada). Slack length was removed, and two fine needle electrodes were positioned to either side of the midline in the belly of the TA. An optimal stimulation amperage was determined via repeated twitch contractions at 0.5 Hz. A tetanic contraction was performed to remove slack from the system before the muscle’s optimal length (L_0_) was determined through the same process. Maximum Tetanic Forces (mN) were normalized to the physiological cross-sectional area (PCSA). The PCSA was determined through the following equation:

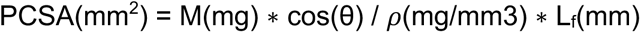

where M is the wet weight of the TA, θ is the pennation angle of the TA, ρ is the physiological density of muscle (1.056 mg/mm^3^), and L_f_ is the fiber length of the TA. L_f_ was converted from the optimal muscle length through the observed fiber length to muscle length ratio of 0.61. To determine the half-max frequency of the force-frequency relationship, the data was fitted to a sigmoidal curve (68). To determine percent force loss as caused by the fatigue protocol, the final tetanic contraction was compared to the starting tetanic contraction.

### Fiber type analysis

Fiber typing followed the methods described in (69). Frozen muscle sections (10µm) of the gastrocnemius muscle were fixed in ice cold acetone for 5 minutes, rinsed in PBS, and then blocked in 1% bovine serum albumin and 10% fetal bovine serum in PBS for 1 hr. For fiber typing, sections were incubated with primary antibodies BA-F8 (1:10), SC-71 (1:30), and BF-F3 (1:10); all from Developmental Studies Hybridoma Bank, Iowa City, IA) overnight at 4°C. Sections were rinsed in PBS plus 1% bovine serum albumin and then incubated with secondary antibodies AlexaFluor350 anti-IgG2b, AlexaFluor488 anti-IgG1, and AlexaFluor594 anti-IgM (A21140, A21121, and 1010111, respectively; Life Technologies) (all used at 1:500) for 1 hr. Following secondary incubation, sections were again rinsed in PBS plus 1% bovine serum albumin. All primary and secondary antibodies were diluted in a 1% bovine serum albumin PBS solution. All steps were carried out at room temperature using room temperature reagents except where noted. All samples were fixed in ProLong® Gold Antifade Mountant (P36930; Thermo Fisher Scientific, Waltham, MA) and imaged on a Keyance microscope. Type 2B, Type 2A, and Type 1 fibers were quantified and expressed as the % of total myofibers per muscle section.

### Cross-sectional area analysis

*Tibilas anterior* (TA) muscles were dissected and frozen in liquid nitrogen. Frozen muscle sections (10µm) of TA muscle were fixed in 4% PFA for 5 minutes, rinsed in PBS, and then blocked in 1% bovine serum albumin and 10% fetal bovine serum in PBS for 1 hr. Anti-laminin-α2 (LAMA2) was used at 1:100 (cat# L0663; Sigma-Aldrich). Secondary Alexa Fluor 488 goat anti-rat (cat# A11006; Invitrogen) was used at 1:2500. Anti-laminin sarcolemmal fluorescence was used to outline individual myofibers to assess individual myofiber cross-sectional area (CSA) automatically using the Matlab program SMASH (70) as described in (69). The entire TA was imaged for analysis using the Keyence microscope fitted with a 10x objective using the Keyence tiling feature.

### RNA isolation, sequencing, and analysis

RNA was isolated from whole quadriceps muscle in the WT and *Sgcg^−/−^*models of the MRL and D2 strains. This was performed in biological triplicate, where 3 female mice 16 weeks of age were used per cohort. TRIzol (71 15596018, Life Technologies) was used to isolate the RNA, which was then filtered with Aurum Total RNA Mini-kit (71 7386820) and suspended into 30ul of RNase-Free water. The RNA samples were indexed and pooled for 100bp single end sequencing. The Illumina RNA Sample Prep Kit version 2.0 generated libraries from these quadriceps and aligned to mm10, following the methods of (72). Quantitation and analysis of transcripts was performed using EdgeR and Counts per million (CPM) were used to calculate differential expression. Heatmaps were generated from Z-scores calculated from CPM gene values.

### Protein isolation

*Quadriceps* muscles were harvested and flash-frozen in liquid nitrogen. Tissues were ground with a mortar and pestle and lysed in Whole Tissue Lysis buffer (50mM HEPES pH 7.5, 150mM NaCl, 2mM EDTA, 10mM NaF, 10mM Na-pyrophosphate, 10% glycerol, 1% Triton X-100, 1mM phenyl-methylsulfonyl fluoride, 1x cOmplete Mini protease inhibitor cocktail [cat # 11836170001; Roche], 1x PhosSTOP [cat# 04906837001; Roche]). The protein concentration was measured with the Quick Start Bradford 1x Dye Reagent (cat# 5000205; Bio-Rad Laboratories).

### Immunoblotting

15μg of protein lysate diluted in 4x Laemmli buffer (cat# 1610747; Bio-Rad Laboratories) was separated on 4-15% Mini-PROTEAN TGX Precast Protein Gels (cat# 4561086; Bio-Rad Laboratories) and transferred to Immun-Blot PVDF Membranes for Protein Blotting (cat# 1620177; Bio-Rad Laboratories). Membranes were blocked using StartingBlock T20 (TBS) Blocking Buffer (cat# 37543; Thermo Scientific). Primary antibodies anti-p-SMAD3 (71 ab52903, Abcam) and anti-SMAD3 (71 ab40854, Abcam) were used at 1:1000. Secondary antibodies conjugated with horseradish peroxidase were used at 1:2,500 (cat# 111035003; Jackson ImmunoResearch Laboratories). ECL Substrate (cat# 1705061; BioRad Laboratories) was briefly applied to the membranes and visualized using an iBright 1500 Imaging System (Invitrogen). Pierce Reversible Protein Stain Kit for PVDF Membranes (including MemCode; cat# 24585; Thermo Fisher Scientific) was used to ensure equal loading. Band intensity was quantified using the gel tool in FIJI (NIH).

### Decellularization

Decellularization was performed as described in (28) with modifications described in (28, 29). Briefly, flash frozen *quadriceps* muscle was cryosectioned at 25μm (Leica CM1950) and mounted on charged Superfrost Plus Microscope Slides (cat# 1255015; Fisher Scientific). Slides were stored at −80°C until decellularization. To decellularize the tissue, slides were thawed at room temperature for 1 hr and then placed in 15mls of 1% UltraPure Sodium Dodecyl Sulfate (SDS) solution (cat# 15553035; Invitrogen) for 5-10 minutes with agitation (40 rpm). Slides were rinsed in PBS (with calcium & magnesium, Cat# 21030CV; Corning) for 45 min followed by 30 min rinse in UltraPure Distilled H_2_O (cat# 10977015; Invitrogen) and then an additional 45 min wash in PBS. Scaffolds were then fixed for immunostaining or used for live cell studies.

### Myoscaffold immunostaining and immunofluorescence imaging

Decellularized muscle cryosections (dECMs) were fixed in 4% paraformaldehyde, rinsed, and blocked with a 1% BSA/10% FBS blocking buffer at room temperature for 1 hour. Scaffolds were incubated in primary antibodies overnight at 4°C at 1:100: anti-laminin-2 (α-2 Chain) (cat# L0663; MilliporeSigma), anti-TGF Beta 1 (cat# 218981AP; Proteintech and cat# MA515065; Invitrogen), anti-Thrombospondin-4 (cat# AF2390; R&D Systems), anti-TGF-β3 (cat# ab15537; Abcam). Secondary antibodies were used at 1:2,500 for 1 hr at room temperature: Alexa Fluor 488 goat anti-rat (cat# A11006; Invitrogen), Alexa Fluor 594 goat anti-rabbit (cat# A11012; Invitrogen) and Alexa Fluor 594 donkey anti-goat (cat# A11058; Invitrogen). The dECMs were mounted with ProLong Gold Antifade Reagent (cat# P36934; Invitrogen). Images were acquired on a Keyence BZ-X810 microscope with identical exposure settings were across genotypes.

### C2C12 differentiation and quantification on dECM

Quadriceps muscles were cryosectioned and decellularized using the dECM method stated above. To reduce variability, scaffolds from all genotypes were generated and seeded in a single batch. Chambers from MatTek Two Well Cell Culture slides (cat# CCS-2; MatTek) were applied to the slides containing the dECM myoscaffolds. The myoscaffolds were incubated for 24hrs in growth media (DMEM (cat# 11995-073; Thermo Scientific) with 10% FBS and 1% Penicillin/Streptomycin (cat# 15070-063; Thermo Scientific). C2C12 myoblasts were seeded onto the myoscaffolds at a density of 100,000 cells in 3mls of growth media. After 24hrs, growth media was removed and replaced with 3mls of differentiation media (DMEM with 2% Horse Serum (cat # 26050088; Thermo Scientific) and 1% Penicillin/Streptomycin). Seeded scaffolds were incubated in differentiation media for 96hrs and subsequently fixed in 4% paraformaldehyde. Scaffolds were blocked for 1 hr in 1% BSA/10% FBS and then co-stained with anti-desmin (1:100; cat# 16520-1-AP; Proteintech) and anti-Laminin-2 (1:100; cat# L0663; Sigma). Alexa Fluor 488 goat anti-rat (cat# A-11006; Invitrogen) and Alexa Fluor 594 goat anti-rabbit (cat# A-11012; Invitrogen) secondaries were used at 1:2500. Scaffolds were imaged identically on the Keyence BX-810 microscope using 10x and 20x objectives. Eighteen images per strain were taken and quantified (obtained from 3 unique animals per strain, 2 scaffolds per animal, 3 images per scaffold). Desmin area, cell morphological parameters, and nuclei count were quantified for each image using the Fiji (NIH).

### Statistical Analyses

Statistical analyses were performed with Prism (GraphPad, La Jolla, CA). When comparing two groups, two-tailed Student’s t-test with Welch’s correction (unequal variances) was used, unless otherwise noted. When comparing three or more groups of data for only one variable, one-way ANOVA with Tukey multi-comparison was used. When comparing data groups for more than one related variable, two-way ANOVA was performed. A p-value less than or equal to 0.05 was considered significant. Error bars represent +/− standard deviation (SD).

## AUTHOR CONTRIBTUITIONS

JO, AL, JK, GL, AD performed experiments. JO, AD and AW evaluated the RNAseq and apatmer analysis. JO and FL completed the muscle mechanics studies and analysis. PP carried out the muscle histology. MH performed mouse husbandry and plethysmography. RC provided critical input in experimental methods. JO, AD, and EM conceived the studies, analyzed the data, and wrote the manuscript.

## ACKNOWLEDGEMENTS

Supported by National Institutes of Health AR052646, NS047726, and HL061322 and Parent Project Muscular Dystrophy.

## Abbreviations

ECM: Extracellular matrix
dECM: decellularized extracellular matrix
LGMD: Limb Girdle Muscular Dystrophy
SDS: sodium dodecyl sulfate
D2: DBA/2J
MRL: Murphy’s Roth Large
TA: *Tibialis anterior*
CSA: cross-sectional area

## SUPPLEMENTAL INFORMATION

**Supplemental Figure 1.**
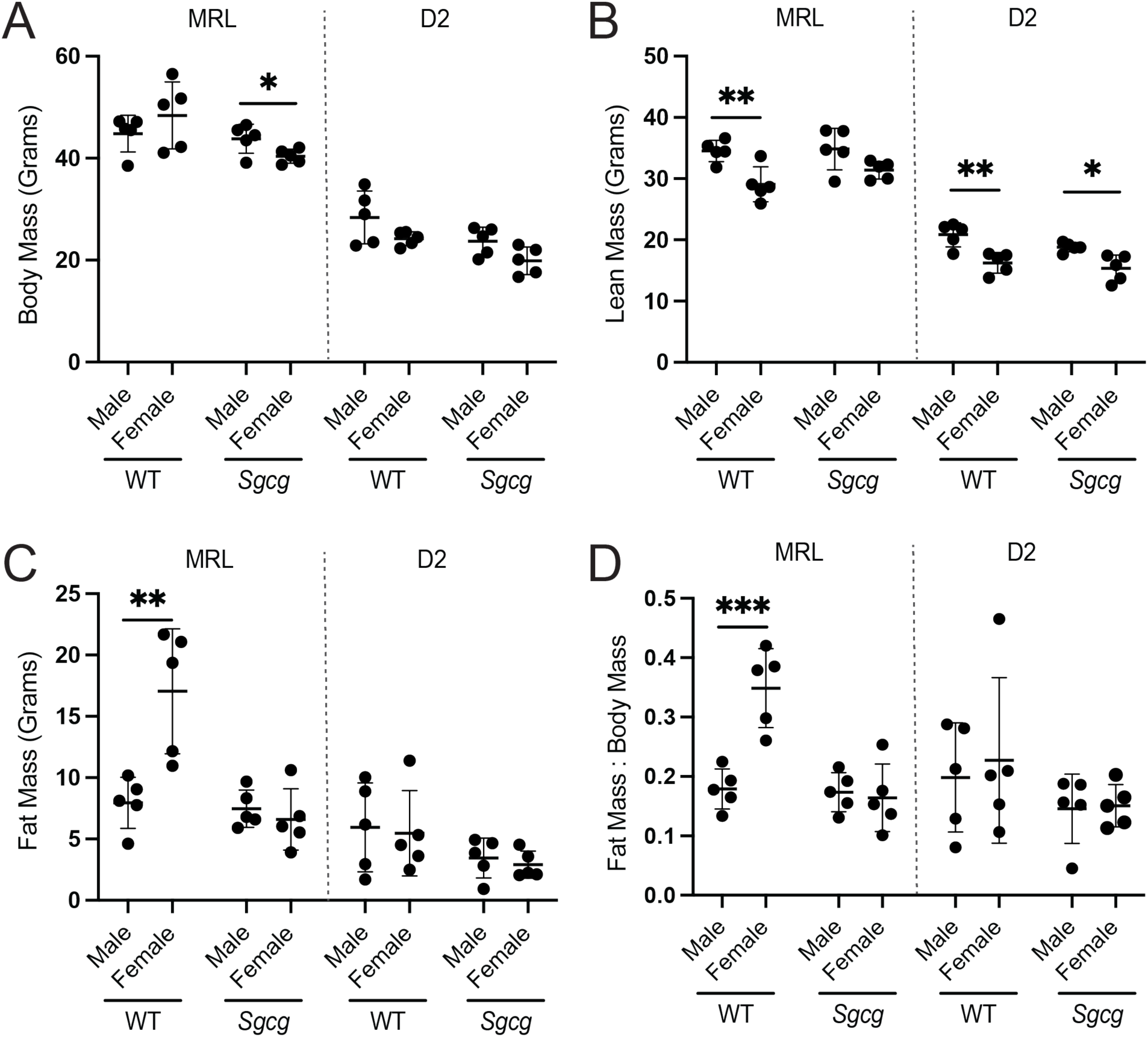
MRL mice were larger than DBA/2J (D2) mice. Whole body mass and body mass composition analysis is shown from 5 male and 5 female mice at 20 weeks. (**A**) Body mass of the MRL strain was significantly higher than the D2 strain, and female *Sgcg*-MRL were lighter than their male counterparts. (**B**) Lean mass was less in female mice compared to male mice, except for *Sgcg*-MRL mice, where this difference was not significant. (**C**) Fat mass of the WT-MRL female MRL mice was significantly greater than WT-MRL male mice. Fat mass remained unchanged between sexes in all other cohorts. (**D**) Fat Mass: Body Mass ratio of the WT-MRL cohort was significantly greater in the females than the males in WT-MRL mice. Graphical quantification of mean ± SD. Student’s t-test was used was used to determine statistical significance. * p<0.05, ** p<0.01, *** p<0.001, **** p<0.0001.

**Supplemental Figure 2.**
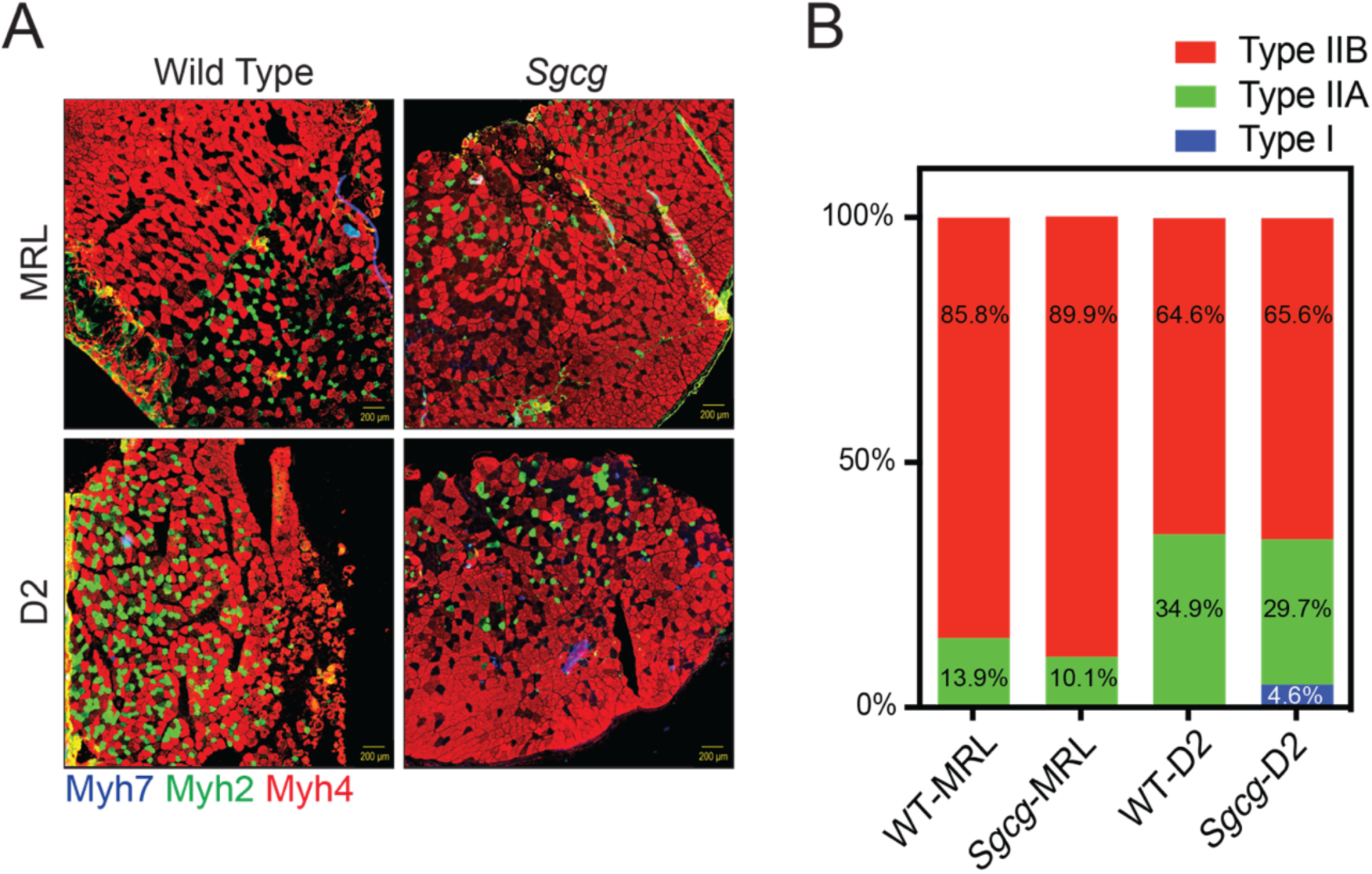
Type I fibers in *Sgcg*-D2 mice were suppressed by the MRL background. *Tibialis anterior* (TA) muscles were harvested from 3 male mice, cryosectioned, and co-stained with antibodies to Myh7, Myh2 and Myh4. (**A**) Representative images of muscles showed a higher proportion of Myh4 positive fibers in the MRL background and a small percentage of Type I fibers in the *Sgcg*-D2 mice. (**B**) Quantified data confirms higher Myh4, lower Myh2 and lower Myh7 positive fibers in the MRL background.

**Supplemental Figure 3.**
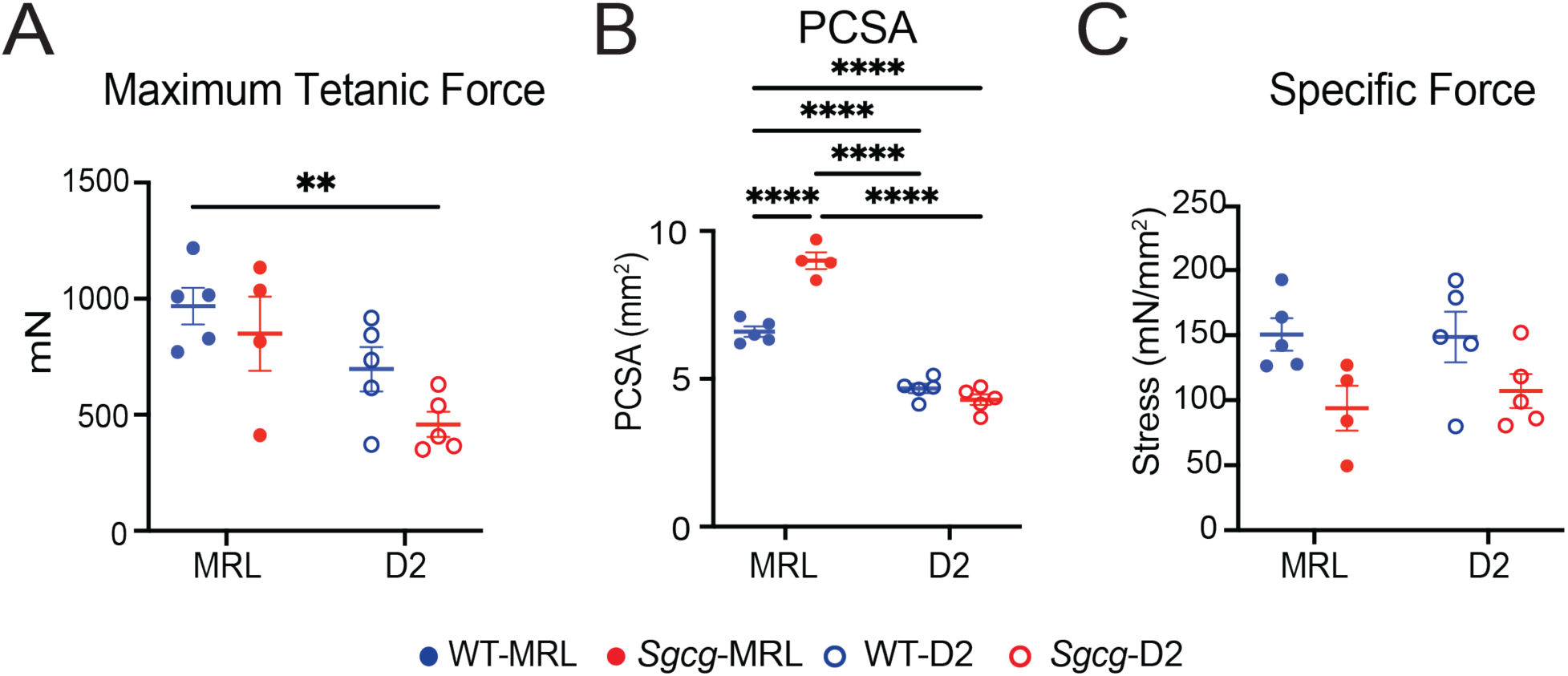
Muscle force measurements demonstrate high variability and no significant differences in force production in *Sgcg* mice from either background strain. Muscle force mechanics were conducted on five female mice, 20 weeks of age. (**A**) The maximum tetanic force in the MRL background trended higher than D2 but did not reach significance. (**B**) Physiological Cross-Sectional Area (PCSA) of the MRL background was significantly larger. **C**. Specific force did not differ by strain or genotype. Graphical quantification of mean ± SD. Two-way ANOVA was used to determine statistical significance. * p<0.05, ** p<0.01, *** p<0.001, **** p< 0.0001.

**Supplemental Figure 4.**
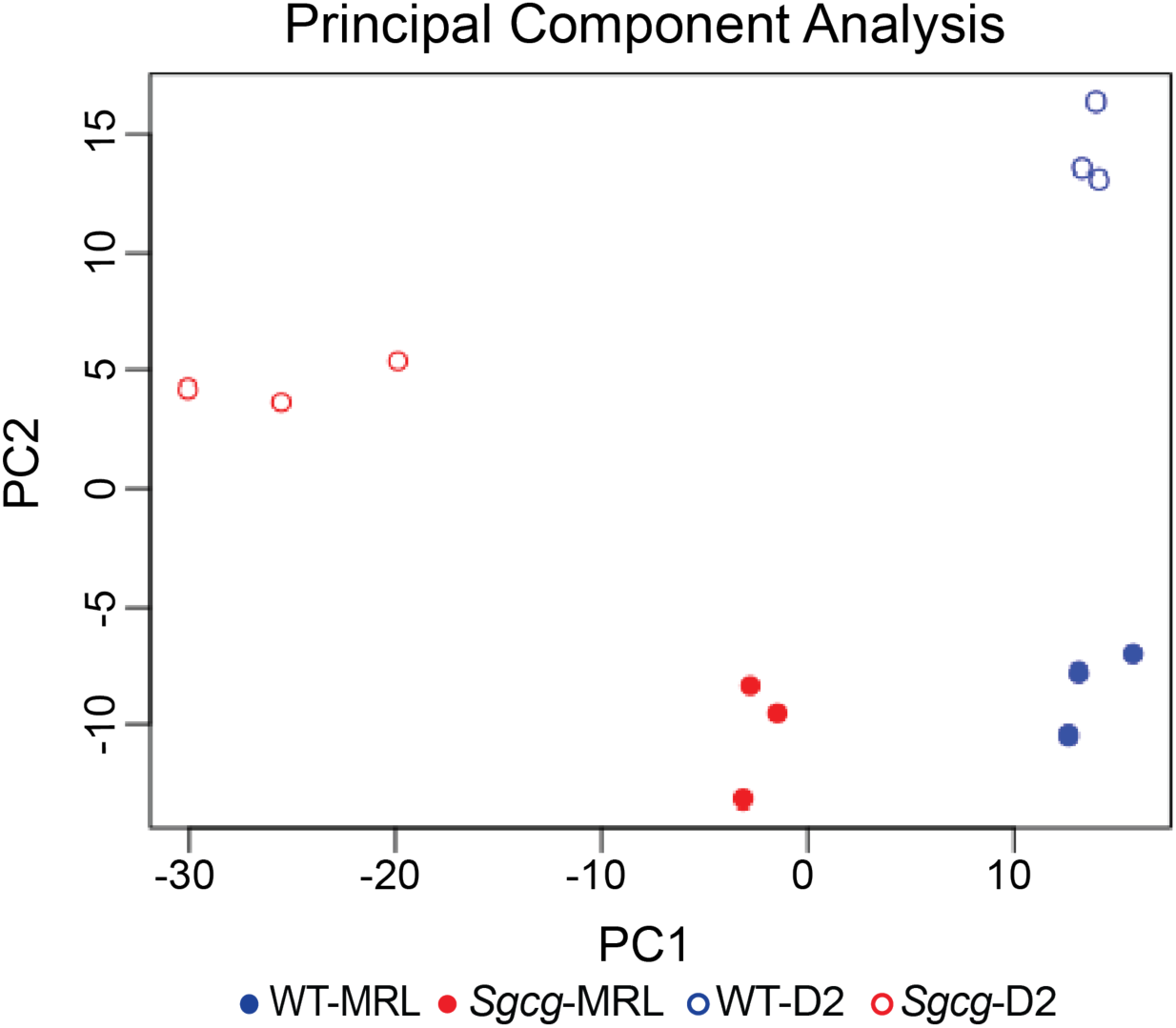
Principal component analysis (PCA) of RNA sequencing shows distinct clustering of phenotypic cohorts.

**Supplemental Figure 5.**
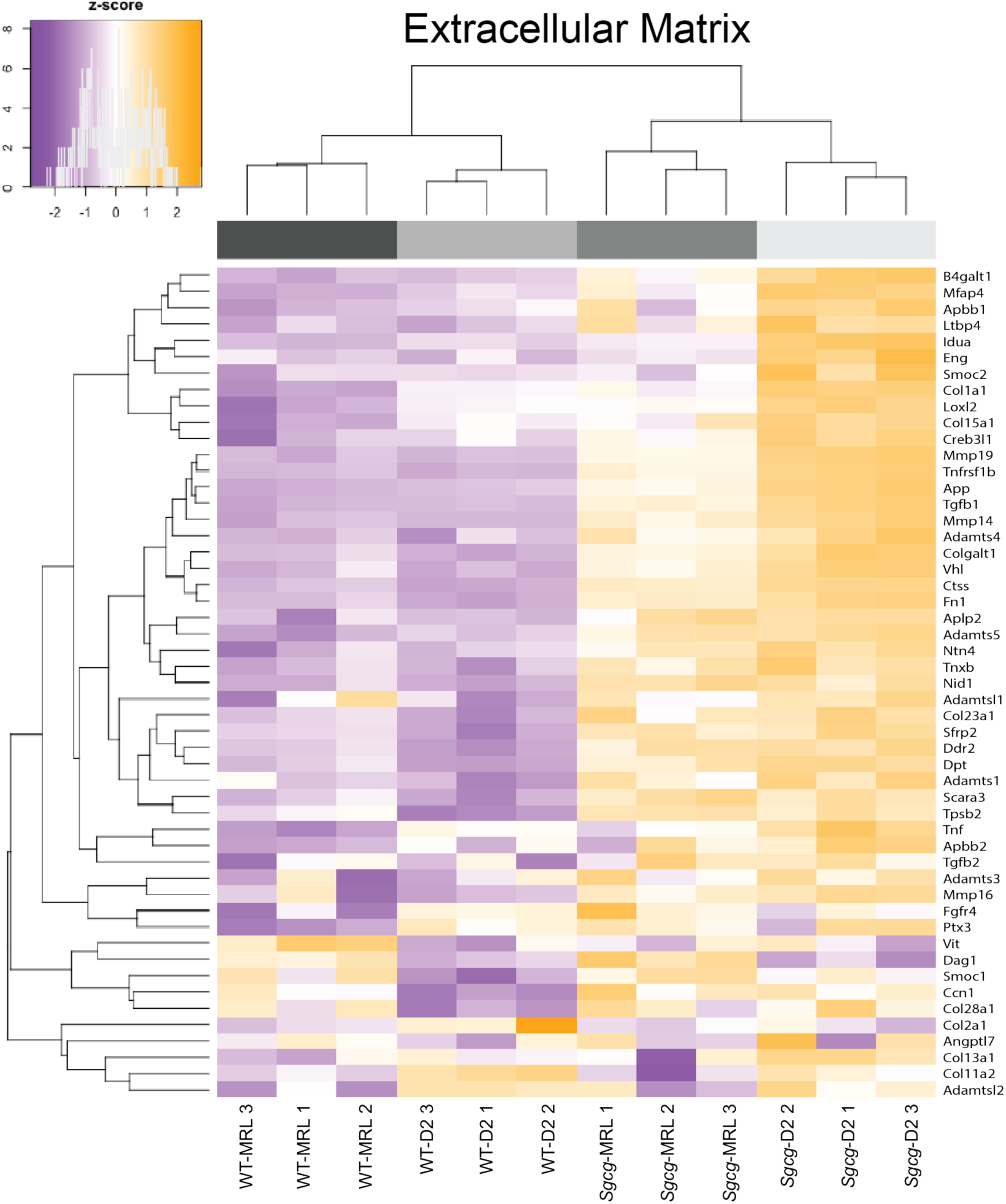
Clustered heatmap indicates upregulation of ECM genes in the *Sgcg*-D2 muscle.

**Supplemental Figure 6.**
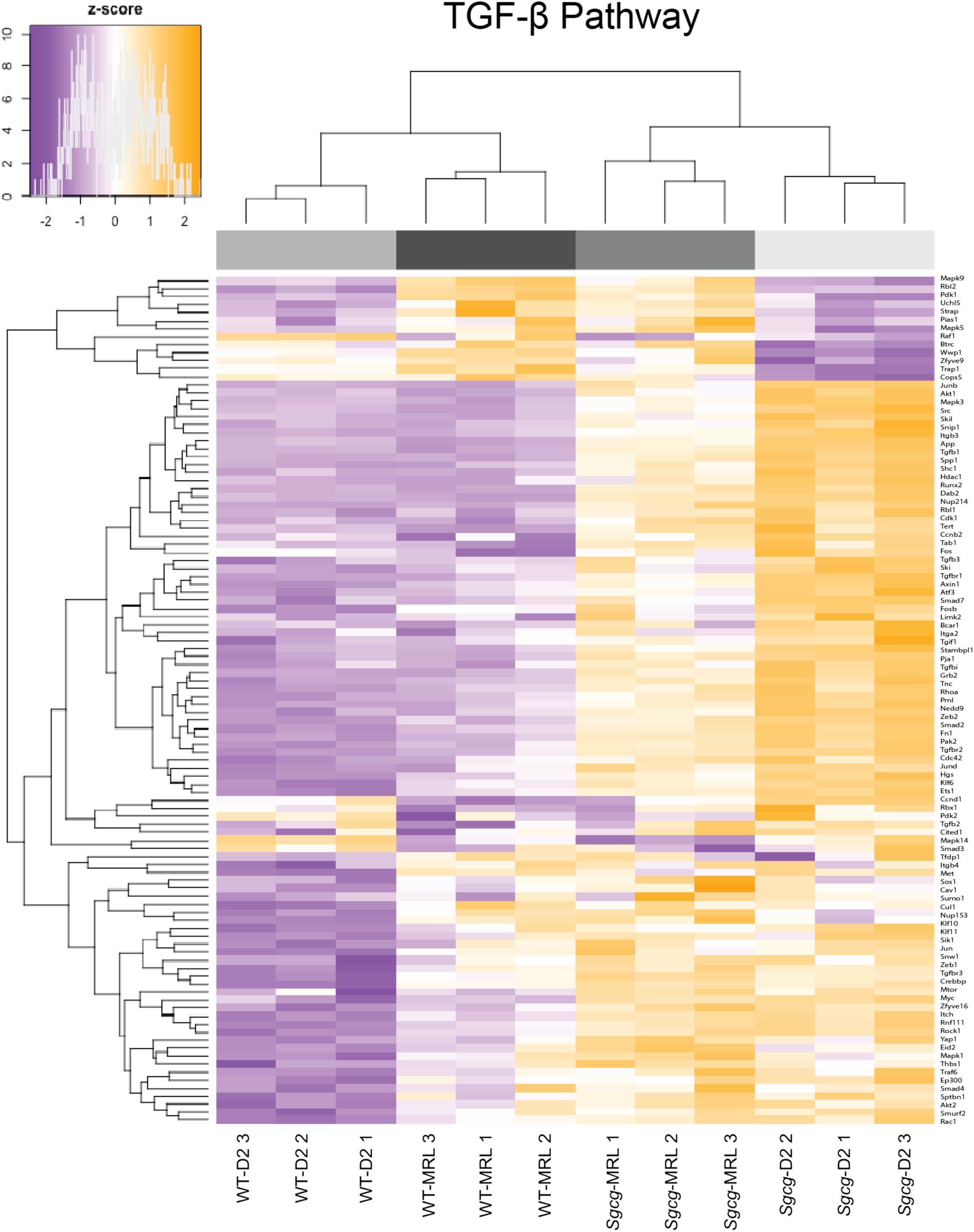
Clustered heatmap illustrating the differential regulation of TGF-β genes in *Sgcg-*MRL compared to *Sgcg*-D2

