## Supplement for "The super-healing MRL strain promotes muscle growth in muscular dystrophy through a regenerative extracellular matrix"

**SUPPLEMENTAL INFORMATION**

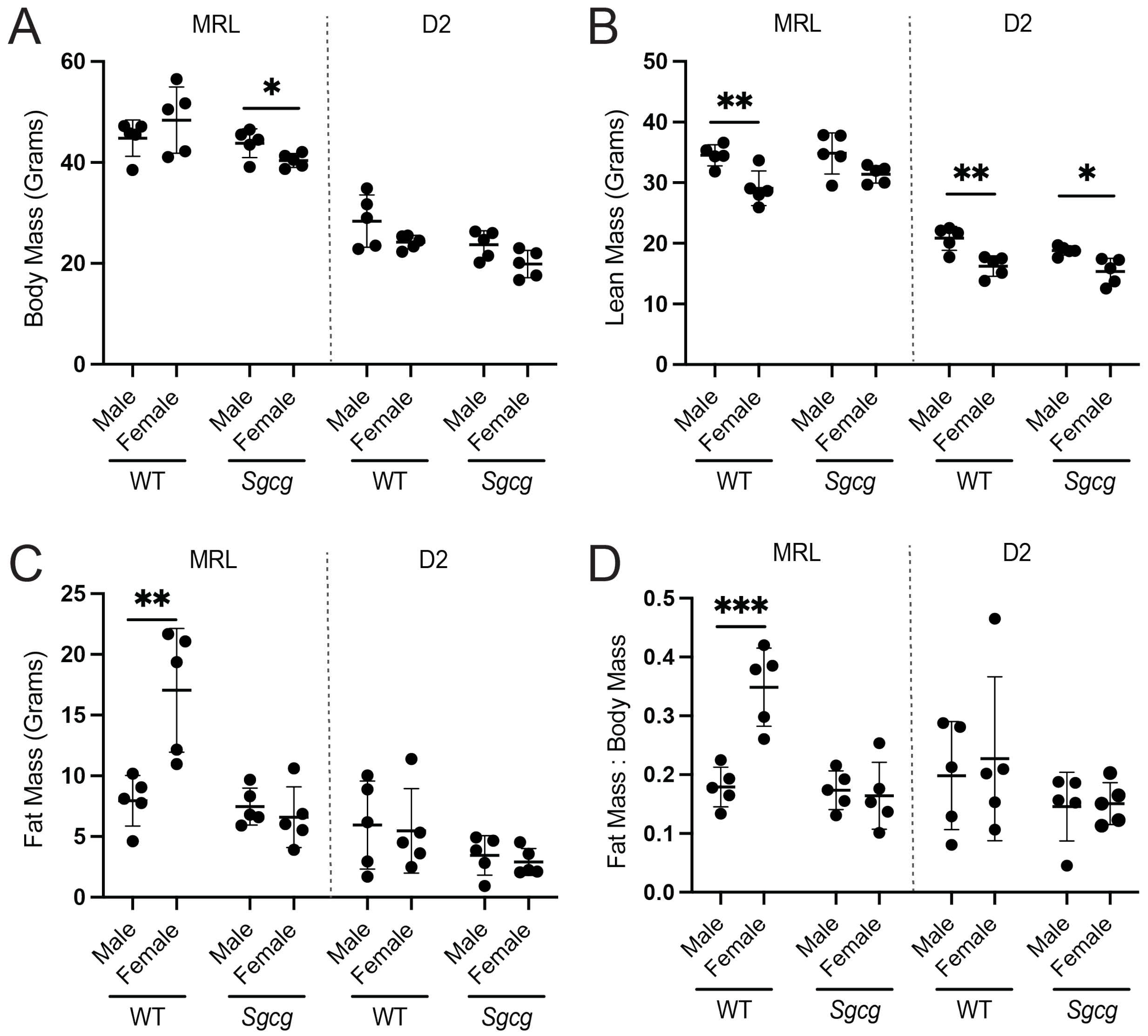


**Supplemental Figure 1. MRL mice were larger than DBA/2J (D2) mice.** Whole body mass and body mass composition analysis is shown from 5 male and 5 female mice at 20 weeks. (**A**) Body mass of the MRL strain was significantly higher than the D2 strain, and female *Sgcg*-MRL were lighter than their male counterparts. (**B**) Lean mass was less in female mice compared to male mice, except for *Sgcg*-MRL mice, where this difference was not significant. (**C**) Fat mass of the WT-MRL female MRL mice was significantly greater than WT-MRL male mice. Fat mass remained unchanged between sexes in all other cohorts. (**D**) Fat Mass: Body Mass ratio of the WT-MRL cohort was significantly greater in the females than the males in WT-MRL mice. Graphical quantification of mean ± SD. Student's t-test was used was used to determine statistical significance. * $p<0.05$, ** $p<0.01$, *** $p<0.001$, **** $p<0.0001$.

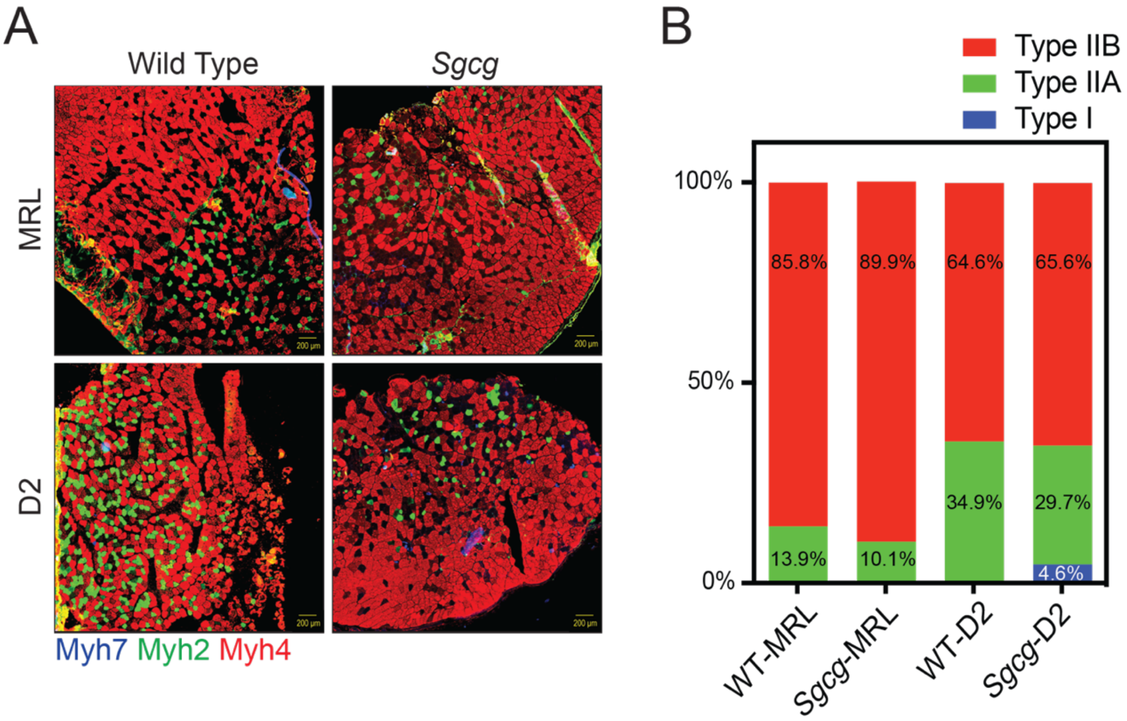


**Supplemental Figure 2. Type I fibers in *Sgcg*-D2 mice were suppressed by the MRL background.** *Tibialis anterior* (TA) muscles were harvested from 3 male mice, cryosectioned, and co-stained with antibodies to Myh7, Myh2 and Myh4. (**A**) Representative images of muscles showed a higher proportion of Myh4 positive fibers in the MRL background and a small percentage of Type I fibers in the *Sgcg*-D2 mice. (**B**) Quantified data confirms higher Myh4, lower Myh2 and lower Myh7 positive fibers in the MRL background.

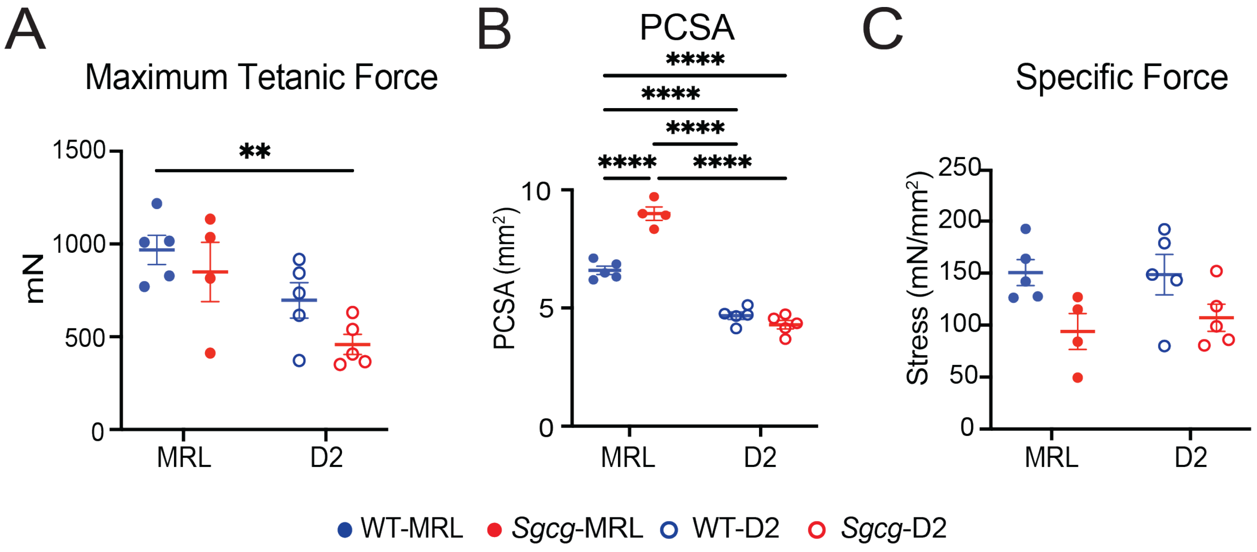


**Supplemental Figure 3. Muscle force measurements demonstrate high variability and no significant differences in force production in *Sgcg* mice from either background strain.** Muscle force mechanics were conducted on five female mice, 20 weeks of age. (**A**) The maximum tetanic force in the MRL background trended higher than D2 but did not reach significance. (**B**) Physiological Cross-Sectional Area (PCSA) of the MRL background was significantly larger. **C**. Specific force did not differ by strain or genotype. Graphical quantification of mean ± SD. Two-way ANOVA was used to determine statistical significance. * p<0.05, ** p<0.01, *** p<0.001, **** p< 0.0001.

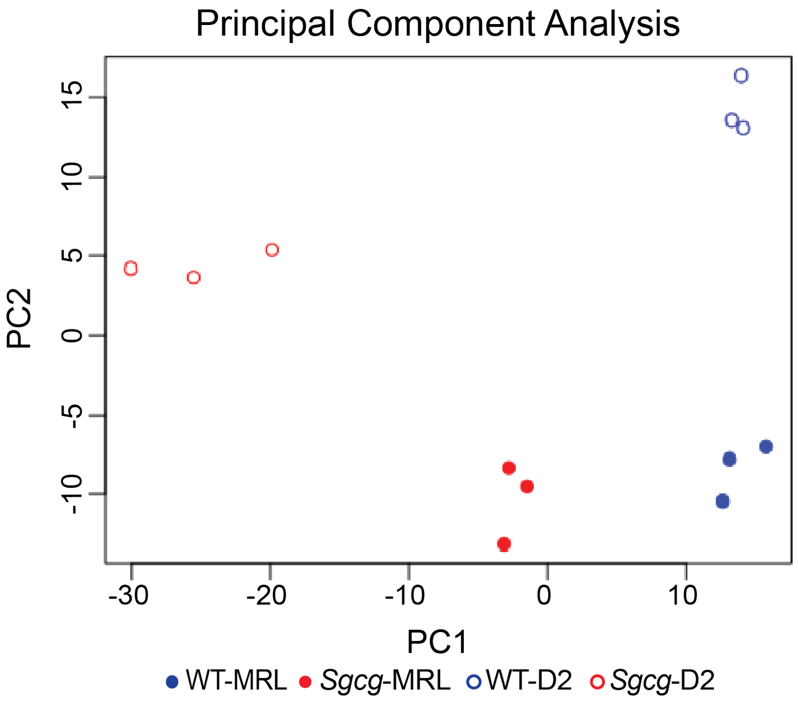


**Supplemental Figure 4.** Principal component analysis (PCA) of RNA sequencing shows distinct clustering of phenotypic cohorts.

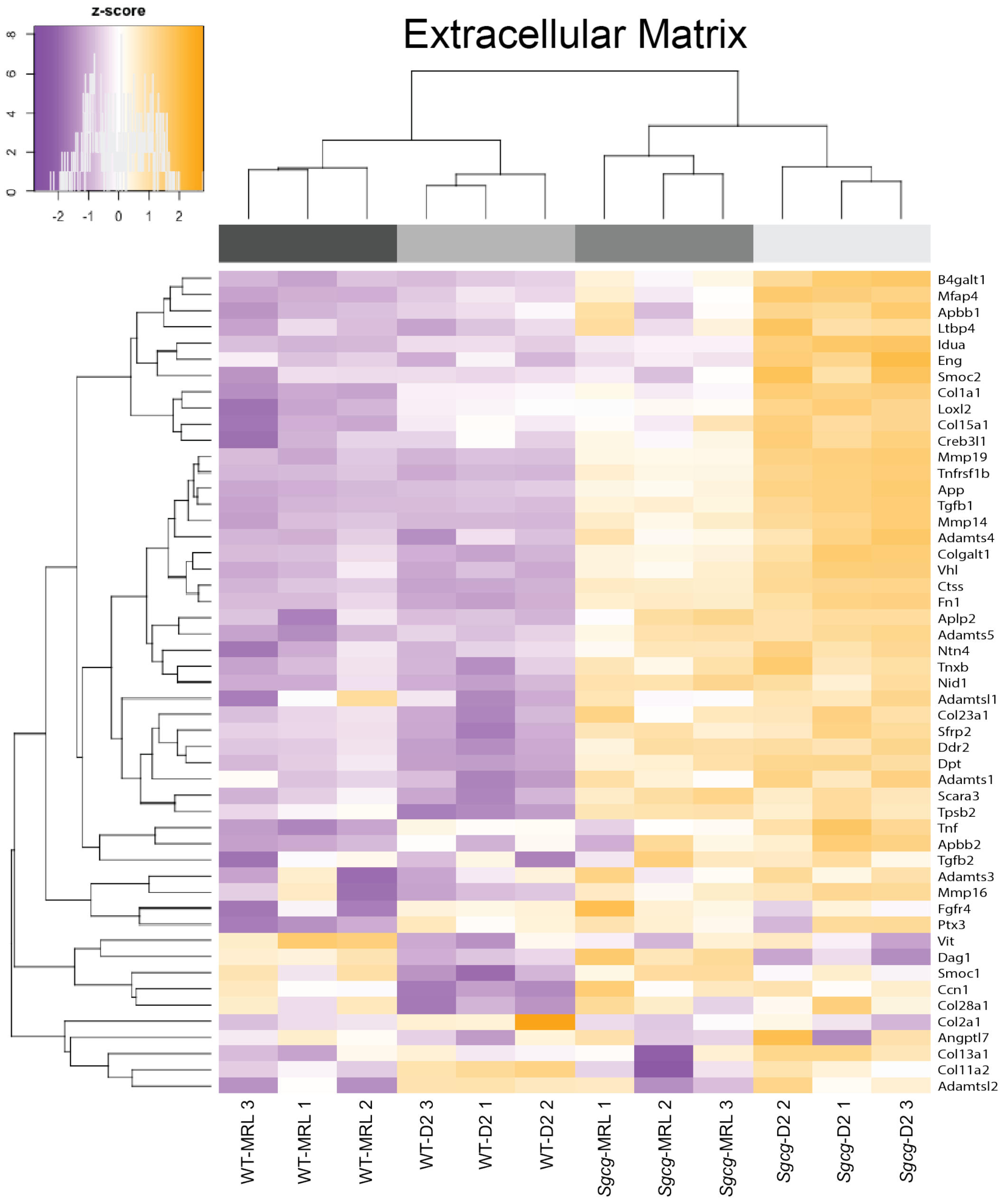


**Supplemental Figure 5.** Clustered heatmap indicates upregulation of ECM genes in the *Sgcg*-D2 muscle.

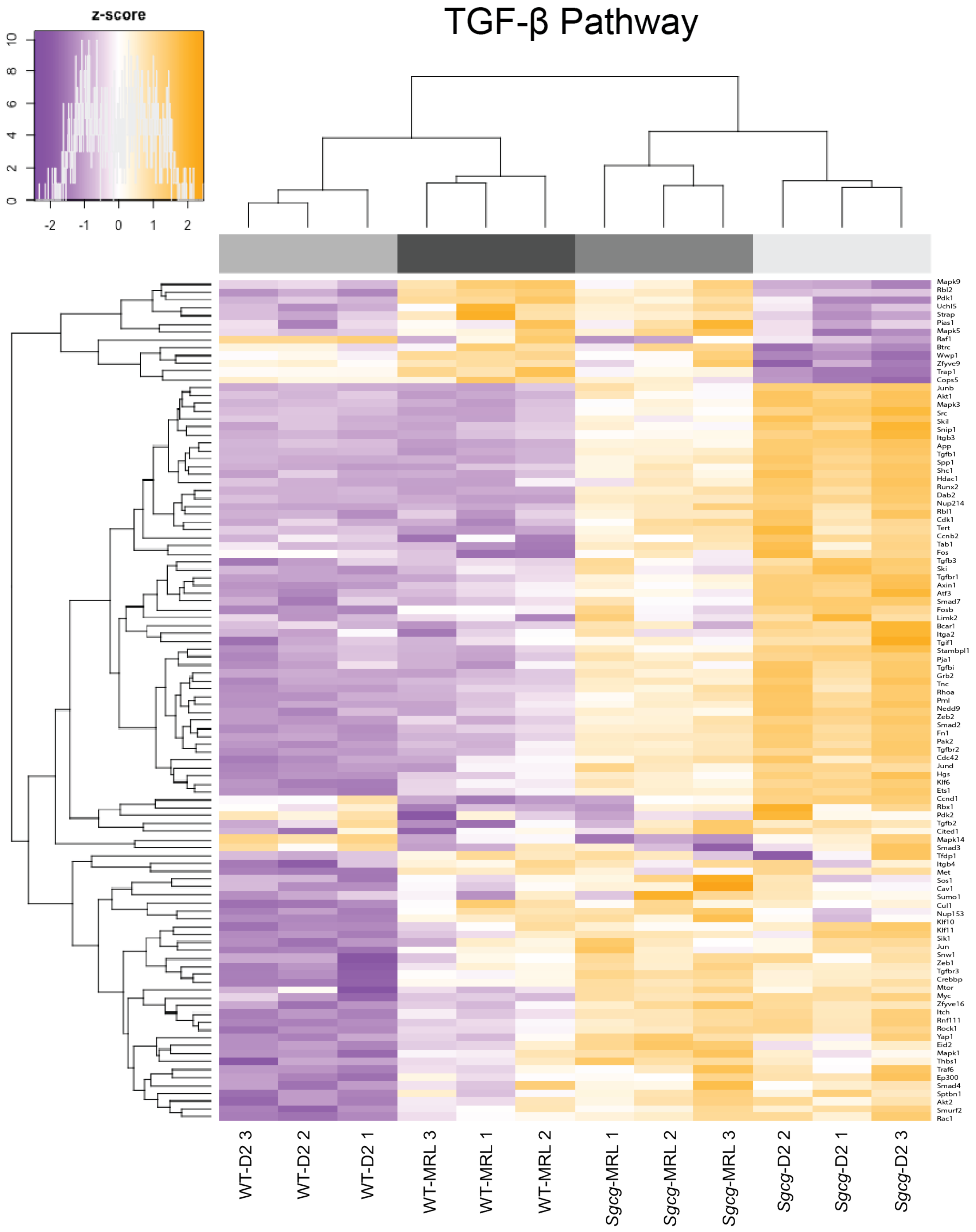


**Supplemental Figure 6.** Clustered heatmap illustrating the differential regulation of TGF-β genes in *Sgcg*-MRL compared to *Sgcg*-D2
